# *Caenorhabditis* diversity on Pohnpei, Micronesia, provides evidence that the Elegans Supergroup has its roots in the Americas and diversified in the Pacific en route to Asia

**DOI:** 10.1101/2025.09.22.677770

**Authors:** Matthew V. Rockman, Sophia C. Tintori, Tuc H. M. Nguyen, V. M. Harmony Yomai

**Affiliations:** Department of Biology and Center for Genomics & Systems Biology, New York University, New York NY; Department of Biological Sciences, Fordham University, Bronx, NY; Department of Fish and Wildlife Conservation, Virginia Tech University, Blacksburg, VA; East-West Center, Honolulu, HI

**Keywords:** *Caenorhabditis*, biogeography, Micronesia, Pohnpei, coexistence, ephemeral patch, dispersal, biodiversity, nematode, population biology, species description

## Abstract

The microscopic nematode *Caenorhabditis elegans* stands unrivaled as a model for developmental biology, neurobiology, and genetics, but fundamental aspects of its ecology, biogeography, and natural history remain unknown. Leveraging recent findings that place its center of diversity in the cool, high-elevation forests of Hawaii, we performed an intensive survey of the *Caenorhabditis* fauna of Pohnpei, a high island in Micronesia that is home to the largest patch of high-elevation forest between Hawaii and East Asia. We found nine species of *Caenorhabditis*, five of them new, but not *C. elegans*. Most species were limited to the hot lowlands but three spanned the elevational range and one was found only in the cloudforest. Using the distribution of *Caenorhabditis* nematodes among habitat patches – individual rotting fruits or flowers – we parameterized simple models that capture key aspects of the population biology of these animals. We generated transcriptomes for the new species and inferred a phylogeny for 70 species of *Caenorhabditis*, based on 2955 genes. This phylogeny allowed us to perform the first quantitative biogeographic analysis for the group. Our analysis suggests that the deep ancestors of the Elegans Supergroup of species lived in the Americas, and that the Supergroup’s subsequent diversification occurred in Remote Oceania. The ancestors of the Supergroup gave rise to a diverse Oceanian fauna and ultimately to multiple lineages that moved into Asia, Africa, Australasia, and back into the Americas. Though biogeographic inferences are limited by the lack of information from key regions of the southwest Pacific, the data are consistent with a model of trans-Pacific migration, with the islands of Oceania serving as sources rather than sinks for biodiversity.

## INTRODUCTION

Over the past half-century, *Caenorhabditis elegans* has emerged as one of our most powerful model organisms (Brenner 1974). Among all the animal species that have ever lived, *C. elegans* is likely the one whose development and neurobiology we understand best. Our exceptional knowledge of these animals’ biology and our power to manipulate them in the lab imbue them with the potential to answer a broad range of fundamental evolutionary and ecological questions (Cutter 2015). Notwithstanding important recent progress (*e.g.,* Devi *et al*. 2025; Crombie *et al*. 2022a; Sloat *et al*. 2022; Ferrari *et al*. 2017; Schulenburg and Félix 2017; Frézal and Félix 2015), however, our knowledge of *Caenorhabditis* ecology and evolution has lagged. Little is known about the local distributions and abundances of individual *Caenorhabditis* species, much less their patterns of cooccurrence and community dynamics.

*Caenorhabditis* has the potential to serve as a powerful model for questions in ecology and evolution. Their profound morphological conservatism (most species are indistinguishable) directs focus to other aspects of their biology that influence population-level processes. In particular, the boom-and-bust demography of these animals – individuals are thought to colonize patches of rotting material, feed on the bacteria there and proliferate, their descendants dispersing to new patches when the resources grow scarce – means that variation in colonization may be as important as variation in resource use. For *Caenorhabditis* to become a model for fundamental questions in ecology and evolution, we need to understand the movement of worms from patch to patch and region to region.

Perhaps the most conspicuous lacuna in our understanding is biogeography: where did *Caenorhabditis* diversify and how did the many species come to occupy their particular locales? Our best model species, *C. elegans,* is globally distributed and locally abundant in many parts of the globe, in Paris and Taipei, Brisbane and Cape Town, San Francisco and Sao Paolo (Lee *et al*. 2021). Recent field studies in the Hawaiian Islands, and associated genomic analyses, have revealed that *C. elegans* diversity is higher within Hawaii than across the whole rest of the globe, with multiple ancient lineages coexisting there (Crombie *et al*. 2019; Lee *et al*. 2021; Crombie *et al*. 2022a); other parts of the planet are home to populations relatively depauperate of genetic diversity, consistent with recent global spread, likely through human commensalism (Andersen *et al*. 2012). Further, Hawaiian *C. elegans* are enriched in native forest and show population genetic patterning consistent with the island progression rule (Shaw and Gillespie 2016), suggesting *in situ* evolution over millions of years (Crombie *et al*. 2022a). In Hawaii, as elsewhere in the tropics, *C. elegans* is restricted to cool, high-elevation habitats.

While population genetics suggests that *C. elegans* has deep history in Hawaii, in the central Pacific, its distantly related sister species, *C. inopinata*, is known only from islands at the western edge of the Pacific—Taiwan and the southern Ryukyus of Japan (Kanzaki *et al*. 2018; Woodruff and Phillips 2018). Successive outgroups to *elegans* and *inopinata* are found either in Taiwan and East Asia or in Hawaii (Fusca *et al*. 2025; Salome-Correa *et al*. 2025), leaving equivocal the locations of their common ancestors and the direction of dispersal across the West Pacific. Several further outgroups are primarily endemic to the Neotropics, suggesting that, somehow, the deep ancestors of *C. elegans* departed from the Americas and left descendants scattered across the central and western Pacific.

To constrain the possible biogeographic histories of *C. elegans* and its relatives in the Elegans Supergroup, and to establish some fundamental characteristics of Caenorhabditis population biology, we characterized the previously unknown *Caenorhabditis* fauna of a locality uniquely suited to inform our understanding: Pohnpei, Micronesia, the higher of the two islands between Hawaii and the western edge of the Pacific that harbor cool, high-elevation forests.

Pohnpei is an isolated volcanic island, part of the Caroline Islands in the Federated States of Micronesia (Figure 1). Pohnpei is located approximately 5,000 km southwest of Honolulu, 1,300 km northeast of the closest islands of Papua New Guinea, and 1,500 km southeast of Guam. The island is 344 square kilometers, and the interior of the island rises to greater than 700 m and is clothed in intact native rainforest and cloudforest (Balick *et al*. 2009). In the vast region bounded by Hawaii to the east, the Philippines, Taiwan, and Japan to the west, and New Guinea and the Solomon Islands to the south, no island reaches 1000 m elevation and only a handful reach 500 m. Pohnpei is by far the largest of these. The others include Kosrae, 500 km southeast of Pohnpei, which has cloudforest like Pohnpei; small, young, uninhabited islands at the north end of the Marianas, some of them active volcanos; and two tiny uninhabited islands in Japan’s Volcano Islands, north of the Marianas. In contrast, the bounding land masses have enormous elevation ranges, with peaks exceeding 4000 m in Hawaii and New Guinea and 3500 m in Japan and Taiwan.

**Figure 1.**
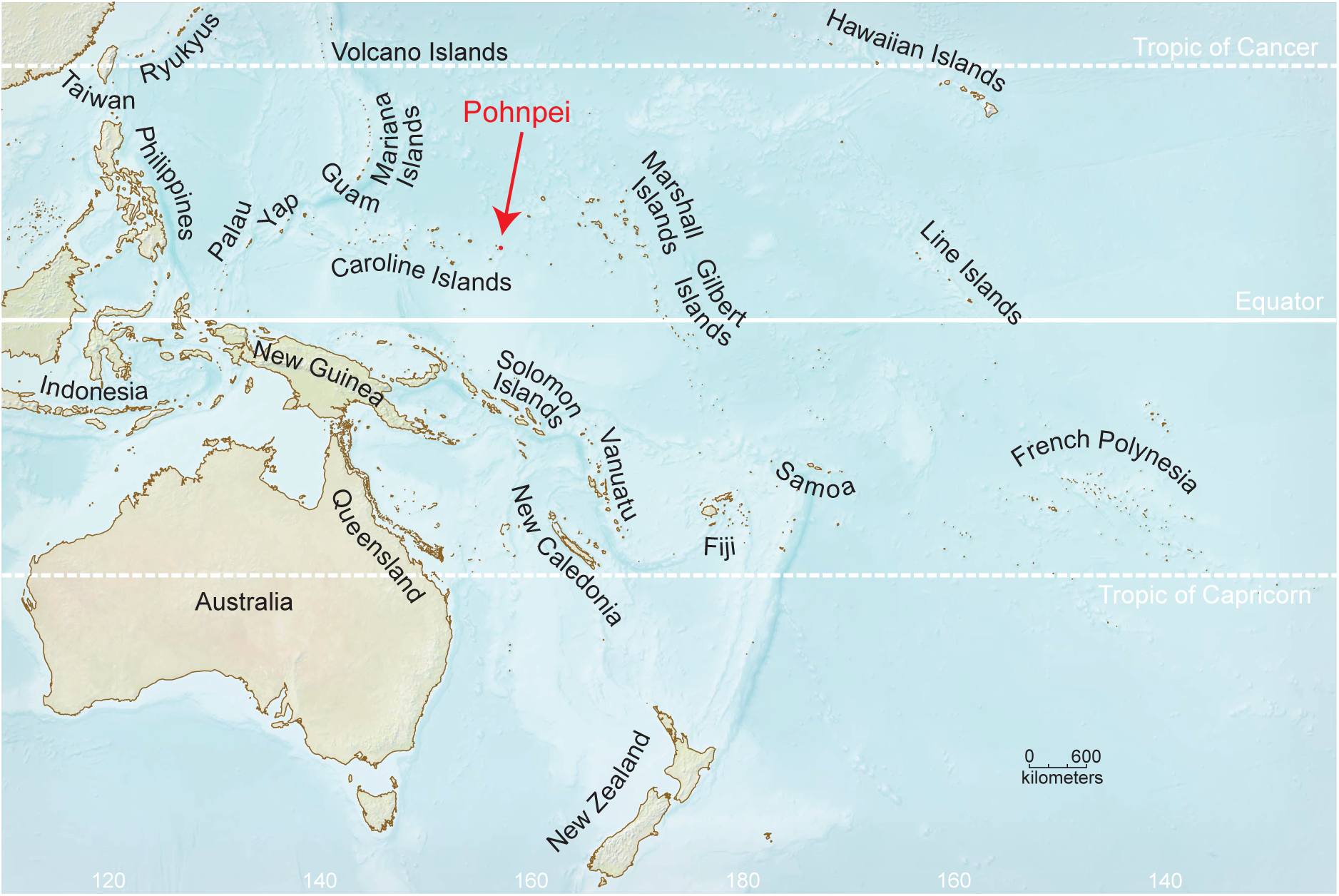
Pohnpei’s location in the Pacific, with small islands, reefs, and atolls magnified. All of the Marshall and Gilbert Islands and most of the Carolines have maximum elevations of just a few meters. This Mercator projection map is adapted from a sourcefile from the University of Texas Perry-Castañeda Library Map Collection.

Located at 7° north of the equator, Pohnpei is hot (mean 27 °C) and exceptionally rainy (4,850 mm/yr) at sea level, while the island’s central mountains are cooler (mean 23 °C at 450m) and even more rainy (> 9,000 mm/yr) (Raynor 1995; Lander and Khosrowpanah 2004). The island is estimated to be approximately 8 million years old (Rehman *et al*. 2013), older than Hawaii’s eight main islands (Price and Clague 2002). For much of the last hundred million years, however, the central and western Pacific were home to a diverse array of high islands, now represented by the guyots and atolls of the Hawaiian-Emperor chain, the Marshall Islands, and the Carolines and Marianas west and northwest of Pohnpei (Nunn 2008).

We used field data, experimental crosses, and *de novo* transcriptome assemblies to discover and characterize the *Caenorhabditis* species of Pohnpei, define aspects of their natural history, and incorporate them into the phylogeny, with implications for the biogeography, ecology, and evolutionary history of this iconic clade.

## METHODS

### Sampling

We collected 400 samples of rotting fruit, flowers, plant stems, fungus, leaf litter, and other potential *Caenorhabditis* habitat patches from eight regions, spanning elevations from sea level to 700 meters and encompassing all of the major forest types on the island, excepting mangroves. Additionally, we collected 111 spatially structured samples from a single locality on Sokehs Ridge. This collection involved three 1 m^2^ quadrats, each gridded into 25 sections, around a nihn fig tree (*Ficus tinctoria*) at 6.9672° N, 158.1907° E.

We recorded collection data using a cell-phone app run on the Fulcrum platform, adapted from the Nematode Field Sampling App (Crombie *et al*. 2022b); this app facilitates the linking of barcoded sample bags to collection metadata. Localities were classified as Urban, Agroforest, Disturbed Forest, Rainforest, or Cloudforest, following Buden (2000), and vegetation samples were identified to species where possible. For each sample, we recorded whether it was collected from the forest floor or from an elevated position (for example, a flower rotting while still on the plant). We measured the surface temperature of each sample in the field using Etekcity Lasergrip 800 temperature guns, and the ambient air temperature using Traceable 4392 Pocket Hygrometer/Dew Point/Thermometer. All localities had relative humidities greater than 70%, and our hygrometer reported humidities of 99.9% most of the time; we therefore have not included humidity as a variable in our analysis.

### Isolating nematodes

Following each day’s collections, samples were placed in Baermann funnels for nematode extractions overnight (Tintori *et al*. 2022). In the morning, nematodes were released onto 6 cm NGM plates seeded with OP50 *E. coli.* Plates were then monitored daily for at least three days for the presence of morphologically typical *Caenorhabditis* nematodes. When we observed candidate *Caenorhabditis* worms, we picked single worms to 3.5 cm plates to establish isofemale or isohermaphrodite cultures. From each survey sample, we aimed to isolate 5 cultures, and for each spatial structure sample we aimed for 10. In many cases, samples contained fewer than the target number of individuals, yielding fewer than 5 or 10 cultures. In some cases, singled worms failed to establish viable cultures (sometimes due to pathogenic bacteria or oomycetes), or candidate worms proved not to be *Caenorhabditis* on further inspection. Every sample determined to have zero *Caenorhabditis* nematodes was verified by a single experienced collector.

### Species identification and cryopreservation

Cultures were transported to New York on their 3.5 cm plates. There, we chunked each culture to a 6 cm plate and allowed it to recover, at which point we bleached it to produce a monoxenic culture on *E. coli* OP50-1 on a 10 cm plate (Stiernagle 2006). When these cultures had expanded to exhaust the food on the 10 cm plate, each was assigned a strain number and cryopreserved.

Many *Caenorhabditis* species are morphologically indistinguishable, and we identified cultures to species using experimental test crosses, following the Biological Species Concept (Félix *et al*. 2014). Worms were maintained at room temperature throughout. For cultures that contained males, we isolated 2 L4 (putative) females to a 6 cm plate and allowed them to mature overnight. If there were no embryos, indicating that the worms were females rather than self-fertile hermaphrodites, we tested whether they belonged to the most common species we recovered, *C. pwilidak sp. nov*, by adding two adult males from an arbitrarily designated reference culture of that species (strain QG4628). Cultures were assigned to *C. pwilidak* if these crosses successfully produced F_2_ larvae; we did not monitor the cultures past that point. In some cases, we observed dead embryos, arrested or deformed larvae, or germline-defective F_1_ adults. In most such cases, a second test cross with QG4628 was more successful, suggesting that the genetically heterogeneous individuals chosen for the crosses varied in their reproductive capacities, but that the cultures were nevertheless conspecific (cultures are isofemale but not isogenic). In some cases, crosses were consistently poor but control crosses between males and females of the focal culture gave similar results, pointing to heritable low fitness under our culture conditions rather than incompatibility with QG4628.

Gonochoristic cultures that failed to produce F_2_s when crossed to QG4628 were crossed to one another and then systematically tested for compatibility, ultimately revealing the presence of six additional gonochoristic species. Each culture’s identity was verified by the production of F_2_ larvae when crossed to the reference culture of one of these other species.

Ten of the 400 survey samples and seven of 111 spatial samples contained gonochoristic *Caenorhabditis* nematodes of a single sex only – one to four females or males – and consequently these could not establish cultures. These individuals were subjected to experimental mating tests with other isolates in Pohnpei, and 20 of the 24 individuals were identified to species by their ability to produce F_1_ and F_2_ descendants from crosses to isolates that were later themselves identified to species in New York.

For androdioecious cultures, we isolated 2 L4 hermaphrodites and then added 2 L4 males of *C. briggsae* QG2801 (AF16 *syIs803, myo-2::GFP*; Inoue *et al*. 2007) and 2 L4 males of *C. tropicalis* QG3501 (NIC58 *qgIs5, myo-3::mCherry*; Noble *et al*. 2021). We then checked the larval offspring of the hermaphrodites for red fluorescence, indicating *C. tropicalis*, or green fluorescence, indicating *C. briggsae*. Cultures were not monitored past the F_1_ larval stage.

The seven gonochoristic species were next characterized by sequencing a fragment of ribosomal DNA, including the *ITS2* region (Kiontke *et al*. 2011). We isolated DNA by DNeasy kit and then PCR-amplified a 2 kb region using primers KK5.8S-1 (ctgcgttacttaccacgaattgcarac) and KK28S-4 (gcggtatttgctactaccayyamgatctgc). We gel-purified the PCR products and Sanger sequenced them (Azenta) with primers KK28S-4, KK28S-10 (gcatagttcaccatctttcgg), and KK28S-22 (cactttcaagcaacccgac). One species (*C. ileile*) did not yield PCR products with any combinations of the primers above, and we generated sequence using primers ST_3242F (cacgaattgcagacgctt) and ST_3951R (ccaaggagcarscgaac). We BLAST-searched the sequences against a database of *Caenorhabditis ITS2* sequences, as well as the NCBI nr database and the Caenorhabditis.org genome database. The newly generated sequences for the 7 gonochoristic species we identified on Pohnpei have been deposited in Genbank with identifiers PP955316-PP955322.

### Ecological statistics

We estimated asymptotic species richness for our 400-sample survey using the *ChaoRichness* function in the iNEXT package (Hsieh *et al*. 2016), with both abundance data (537 individuals) and species incidence data (400 samples).

To infer properties of the local patch colonization process, we make use of several classes of observation. The quantities of interest are the number of colonization events per patch and the number of worms arriving on a patch with each colonization event. Because a successful colonization results in rapid proliferation of the population, these quantities cannot be directly observed. The quantities we observe are the total number of available patches, the number of such patches that have been colonized, and, for gonochoristic (male/female) species, the number of colonized patches that contain only a single worm of a particular species. This last quantity is unavailable for androdioecious species, because a single hermaphrodite can initiate proliferation.

To estimate the number of colonization events per patch, we assume that each patch (a single sampled substrate) receives migrants carried by phoretic vectors – insects or snails, for example – that carry pre-reproductive dauer larvae. Following Sloat *et al*. (2022), we assume that colonization events are independent, and therefore the number of colonization events per patch, *C*, is Poisson distributed with mean *λ_C_*. In a region where a worm species is abundant and there are many equivalent patches available, we can estimate the mean number of successful colonization events per patch using the proportion of patches that contain zero individuals of a species. Given *n* patches, of which *k* have zero worms, the likelihood of colonization rate *λ_C_* is given by 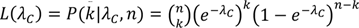. We numerically maximized the likelihood using the *optimize* function in R and estimated the 95% confidence interval from the 1.92-log drop in log likelihood.

To estimate the number of founder dauers delivered at each colonization event, *n_d_,* we assume that dauers behave independently and therefore this number is also Poisson distributed, zero-truncated because we condition on colonization. We estimated *n_d_* for a gonochoristic species, using counts of patches that contain exactly one worm. Such patches occur when there has been exactly one colonization event and it bought exactly one worm. The probability of observing *k* patches that contain exactly one worm of a gonochoristic species, conditional on observing *n* patches that contain at least one worm of that species, can be modeled as a binomial whose success probability *p* is the product of a zero-truncated Poisson probability of a single colonization event, with Poisson parameter *λ_C_*, and a zero-truncated Poisson probability that the number of dauers transferred is 1, given the Poisson parameter *λ_D_*. This works out to 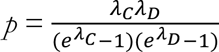, and 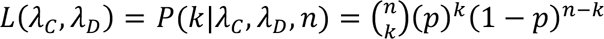. With an estimate of the parameter *λ_D_*, we then find the expected number of dauers per colonization, *n_d_,* as the mean of the zero-truncated Poisson: 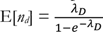. We numerically found the joint likelihood surface for the two lambda parameters, and also used *optimize* to maximize the likelihood of *λ_D_* conditioned on our estimates of *λ_C_*. Accurate counts of one-worm patches requires that the single worms be identified to species, which in our case involves experimental test crosses in the field. Consequently, we have reliable data only for *C. pwilidak* sampled in the 400-sample survey.

Simulations of patch colonization processes using the parameter estimates of 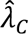 and 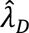 were implemented in R and are presented in File S1.

To test for clustering of worms in our spatial samples at Sokehs Ridge, we calculated Geary’s (1954) *c*. Physical distances ranged between 0.2 m (for adjacent sections within quadrats) and 5 m (for different quadrats). Worm densities were quantified with a rank score between 0 and 6 (representing 0 worms, 1-5, 6-10, 11-50, 51-100, 101-500, and 501-1000, respectively). For species-level analyses, we assigned each sample a value of 0 or 1 (representing absence or presence, respectively) and calculated Geary’s *c* using this attribute.

### Transcriptome sequencing and assembly

To generate data for phylogenetic analysis, we performed RNAseq on populations of worms treated to maximize transcriptomic diversity. For each species-reference isofemale line, we pooled mixed-sex, mixed-stage worms from five different growth conditions, as in Sloat *et al*. (2022): one population was fed with bacteria from the *Ce*MBio collection of strains that mimics the *C. elegans* diet (Dirksen *et al*. 2020), and the other four were maintained on a standard laboratory diet (*E. coli* OP50) but then subjected to either starvation, heat stress, cold stress, or standard well-fed conditions. Temperature stress consisted of exposing the worms to either 35° C or 4° C for 2 hours followed by a 2-hour recovery.

Pools of worms were flash-frozen in liquid nitrogen, and total RNA was extracted using a trizol+chloroform protocol and an RNAeasy spin-column cleanup including a DNAse treatment. We then used the Illumina Stranded mRNA Prep kit to generate mRNA, convert it to cDNA, ligate adapters from the IDT for Illumina RNA UD indices set B, and amplify the library. The library was sequenced on NextSeq500 with a MidOutput 2×150 v2.5 configuration at NYU’s GenCore facility. The GenCore staff used Picard IlluminaBasecallsToFastq version 2.23.8 (https://broadinstitute.github.io/picard/) for basecalling, with APPLY_EAMSS_FILTER set to false, and Pheniqs version 2.1.0 (Galanti *et al*. 2021) for demultiplexing.

Following the protocol outlined in Sloat *et al*. (2022), we first trimmed adapters from the paired-end sequences using TrimGalore (https://github.com/FelixKrueger/TrimGalore). Trimmed sequences were assembled into transcriptomes using Trinity 2.15.1 (Grabherr *et al*. 2011), running default parameters for paired-end reads. The RNAseq data (paired-end fastqs) and transcriptome assemblies for the five new Pohnpeian species are reported in BioProject ID PRJNA1128046.

### Phylogenetic inference

We used TransDecoder 5.5.0 (https://github.com/TransDecoder/TransDecoder) to extract protein sequences for the longest predicted ORF for each transcript from our transcriptome assemblies. We collected analogous protein sequence FASTA files for an additional 65 *Caenorhabditis* species, as detailed in Table S1; the bulk of the data are drawn from the *Caenorhabditis* Genomes Project (Stevens 2024).

To isolate sets of homologous genes from each species, we applied BUSCO 5.3.0 (Seppey *et al*. 2019), with the nematode_odb10 dataset, to each of the protein FASTA files. For each protein sequence in its database, BUSCO identifies homologous sequences in the input file and classifies them as single or multicopy within each species. The transcriptomes of our new Pohnpeian species had a greater fraction of inferred multicopy genes than is typical for the other input files, including our own transcriptomes for other species (Table S1). This pattern may reflect the fact that the Pohnpeian transcriptomes were generated from isofemale lines, which are segregating genetic variation from a sample of four genomes (two diploid parents), while the other samples are derived from fully or largely homozygous inbred lines; that is, inferred multicopy genes may represent alternate alleles of single-copy genes.

Following the “ONE” approach outlined in Yan *et al*. (2022), we generated a dataset for each BUSCO gene incorporating data from each species in which the gene was classified as “complete.” These datasets include single-copy genes from some species; in other species, which are inferred to carry multiple paralogs, we chose the longest ORF from the set of multiple genes classified as belonging to that BUSCO gene family. In many cases the paralogs within a species likely represent species-specific duplicates or alleles, in which case the relationship between species tree and gene tree is unaffected by paralogy. More generally, Yan *et al*. (2022) provide justification and validation for the use of paralogs in species-tree inference under the multi-species coalescent. In the present case, inclusion of paralogs dramatically increases the representation of data for each species and the total number of genes for analysis.

Next, we reduced the dataset to BUSCO gene families with data from at least 80% of the 70 species, and we used MAFFT 7.475 (Katoh and Standley 2013) to align the sequences, with default settings. We eliminated poorly aligned regions using TrimAl 1.4.1 (Capella-Gutierrez *et al*. 2009) with settings -gt 0.8 -st 0.001 -resoverlap 0.75 -seqoverlap 80. This process yielded trimmed alignments for 2,955 genes. We estimated maximum-likelihood gene trees using IQ-TREE (Nguyen *et al*. 2015) with the LG+I+G model (Yang 1994; Le and Gascuel 2008). We then estimated a species tree from the collection of gene trees under the coalescent model implemented in ASTRAL-III 5.7.8 (Zhang *et al*. 2018), and we bootstrapped over gene trees to estimate branch support.

To estimate branch lengths, we again followed Yan *et al*. (2022) and used sequences from inferred single-copy orthologs only. The dataset includes 720,649 amino acid positions from 1,913 BUSCO genes present in single-copy in at least 80% of the species. We concatenated the sequence alignments using catfasta2phyml (https://github.com/nylander/catfasta2phyml) and estimated branch lengths for the 2,955-gene species tree using this concatenated dataset.

### Biogeography

Biogeographic data derive from sources cited in Table S2. *Caenorhabditis* researchers share information via several informal channels, and we drew heavily from strain data in RhabditinaDB v. 0.92 (Fitch *et al*. 2024) and collection data in evolution.wormbase.org, a community wiki resource.

To reconstruct ancestral ranges, we used maximum likelihood inference implemented in BioGeoBEARS 1.1.3 (Matzke 2018). We restricted analysis to the Elegans Supergroup and its three successive outgroups (the Guadeloupensis Group, *C. astrocarya*, and the Drosophilae Supergroup). The more distant lineages that we excluded (7 species) are sparsely sampled and have heterogeneous branch lengths. We rendered the remaining phylogeny ultrametric using the correlated rates model implemented in *chronos* function in the ape package v.5.7-1 (Sanderson 2002; Paradis 2013; Paradis and Schliep 2019).

With a focus on the largest-scale of biogeographic history, we partitioned the globe into five broad regions. Four of these represent continents: the Americas, Africa, Eurasia (including western Indonesia, Japan, and Taiwan), and Australasia (represented by data from Australia and the islands of Malaita and Guadalcanal in the Solomon Islands). The fifth region is Remote Oceania, represented by data from Pohnpei, the Hawaiian Islands, and Moorea in French Polynesia. The precise boundaries of these regions are relatively unimportant because most of the boundary areas have never been sampled. For example, we have no information from Vanuatu, Fiji, Samoa, the Philippines, or New Guinea, and small-scale sampling in New Caledonia and New Zealand has uncovered only the globally distributed species *C. briggsae* and *C. elegans*. For the regional assignments with some ambiguity, we let previous biogeographic work, and our phylogenetic results, guide our decisions (Bernstein *et al*. 2023; Holzmeyer *et al*. 2023). We assigned the Solomon Islands to Australasia because of their botanical affinities and because the Islands were nearly connected to New Guinea and Australia during the Pleistocene glacial periods (Keppel *et al*. 2009; Holzmeyer *et al*. 2023). In Indonesia, data come only from a small radius encompassing East Java, Bali, Lombok, and the nearest corner of Sulawesi (Devi *et al*. 2025). Because Java and Bali were connected to Asia as part of Sundaland during the Pleistocene, and because four of the *Caenorhabditis* species found on Java and Bali were also found on Sulawesi and Lombok, we assigned the whole region to Asia. Finally, we treated the isolated islands of Remote Oceania as a single biogeographic region, because the species from Hawaii, Pohnpei, and French Polynesia are tightly clustered in our phylogeny. Overall, we view our biogeographic analysis as a crude first look, with denser sampling of critical western Pacific regions as necessary next step.

Most of the species with widespread distributions encompassing distant continents are inferred to have achieved these distributions recently, through anthropogenic activity, and their pre-human distributions are unknown. We therefore excluded these species from analysis (*C. sp. 8, C. remanei, C. nigoni, C. brenneri, C. briggsae, C. tropicalis,* and *C. zanzibari*). We made two exceptions to this approach: we retained *C. elegans* and coded it as having a Remote Oceania range, based on the population genetic data linking it to Hawaii (Andersen *et al*. 2012; Crombie *et al*. 2019; Lee *et al*. 2021; Crombie *et al*. 2022a), and we retained *C. imperialis*, known from both French Polynesia and the Lesser Antilles, and we coded it as Remote Oceania exclusively, based on its close relationship to other exclusively Remote Oceania species. We also excluded *C. sp. 2*, which is known only from the Mediterranean, Madeira, and Canary Islands, but exclusively in association with *Opuntia* cactus, a recent introduction from the New World (Kiontke and Sudhaus 2006). We performed inference on the resulting 55-species phylogeny. We fit a Dispersal-Extinction-Cladogenesis (DEC) model (Ree and Smith 2008) and two other standard biogeographic models (Matzke 2018), as well as each of these models incorporating an additional parameter (+J) allowing simultaneous dispersal and speciation (Matzke 2014; Revell and Harmon 2022).

To estimate ages for nodes on the phylogeny, we used the branch lengths from the time tree of Fusca *et al*. (2025), who performed a Bayesian analysis that used laboratory-based per-generation mutation-rate estimates (Saxena *et al*. 2019) to date nodes in units of generations, after carefully accounting for synonymous site saturation and codon usage bias. Those branch lengths were highly correlated with those we estimated for our phylogeny (adjusted R^2^ = 0.98). We regressed the branch lengths from our ultrametric biogeography tree onto the matched branches from the timetree, and used the slope to rescale all of the branches of the biogeography tree in units of generations.

### Species descriptions

Descriptions and nomenclatural acts are presented in Appendix 1. Type cultures of each new species have been deposited with the *Caenorhabditis* Genetics Center.

## RESULTS

### Species diversity and natural history

We collected 400 samples of putative *Caenorhabditis* habitat – rotting fruit, flowers, fungi, and vegetal litter, primarily – from all five municipalities of Pohnpei (Figure 2A, Table S3). We sampled agroforest, rainforest, and cloudforest, as well as some sites in disturbed forest and urban settings.

**Figure 2.**
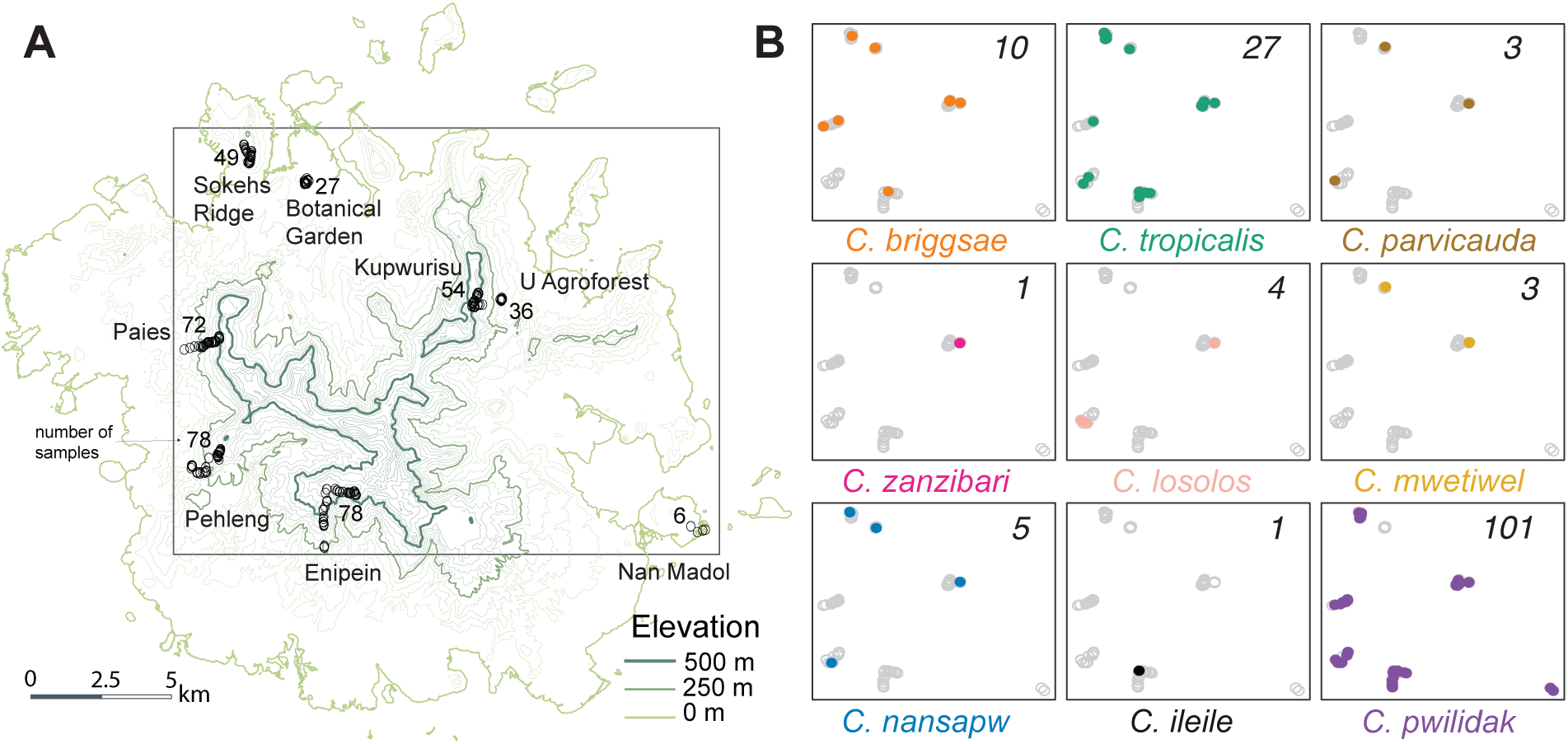
A. We collected 400 samples from eight regions of Pohnpei, at elevations from sea level to 705 m. **B.** Colored dots indicate the presence of each of the 9 *Caenorhabditis* species across the sampled regions. Each box plots the boxed region of panel A, and the numbers report the number of samples that contained each species.

We used Baermann funnels (Tintori *et al*. 2022) to extract the nematodes from each sample. In total, 88% (351) of the samples yielded nematodes, and 35% (140 samples) yielded *Caenorhabditis*.

From the 140 samples containing *Caenorhabditis*, we identified 537 individual worms (Table S4) to species (five *Caenorhabditis* individuals per sample when possible). These worms represent nine species of *Caenorhabditis*, mutually incompatible with each other (no F_2_s) in experimental test crosses. The point estimate of asymptotic species richness, based on both individual and sample rarefaction, is 10, suggesting that our survey has nearly saturated the species diversity given our sampling strategy (Chao *et al*. 2014; Hsieh *et al*. 2016). The species collected include *C. parvicauda, C. zanzibari, C. briggsae, C. tropicalis*, and five new species that are dissimilar to all known species in ribosomal DNA sequences and reproductively incompatible with the known species that share the greatest sequence similarity (Table S5). These species are formally described and named in Appendix 1. *C. briggsae* and *C. tropicalis* are androdioecious species (males and self-fertile hermaphrodites); the other seven species are gonochoristic (males and females).

*C. pwilidak sp. nov.* was the most abundant and widespread species (present in 101 samples), followed by *C. tropicalis* (27 samples) and *C. briggsae* (10 samples). Two species, *C. ileile sp. nov.* and *C. zanzibari*, were each observed only once, in each case co-occurring with other species. All other species were found in multiple localities on opposite sides of Pohnpei’s central mountains (Figure 2B). Co-occurrence of multiple species within individual samples was common (Figure 3A).

**Figure 3.**
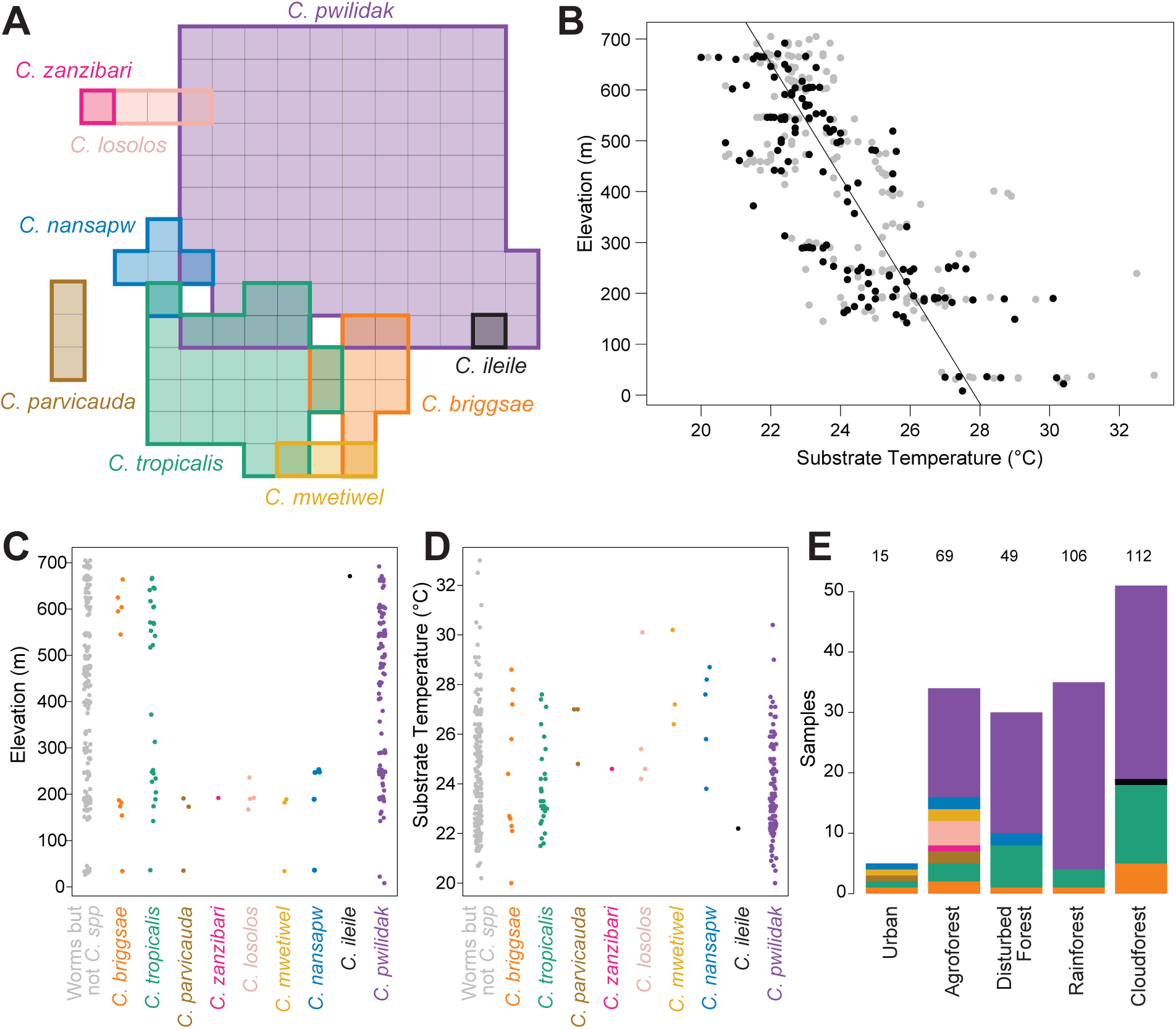
**A**. Eight of the nine species co-occurred with other *Caenorhabditis* species in individual samples. Here, each box represents a sample that contained at least one *Caenorhabditis* species. **B.** Substrate temperatures declined with elevation. Gray dots represent samples that contained worms, and black dots are the subset that contained *Caenorhabditis*. The line shows a linear fit of temperature on elevation for all worm-containing substrates. **C.** Species distributions as a function of elevation, and **D**, as a function of temperature. **E.** Counts of samples containing each species for each forest type, ordered by mean elevation. Colors represent species, as in panels A, C, and D. Numbers at the top indicate the total number of worm-containing samples collected in each forest type.

Field-measured substrate temperatures showed the expected relationship with elevation (Figure 3B), dropping 0.9°C with each 100 meters. Many of the variables that likely influence species distributions – elevation, temperature, rainfall, substrate types, and forest types – are highly correlated on Pohnpei, reducing our ability to make strong statistical inferences. Nevertheless, there are some suggestive patterns (Figure 3C-E).

The three common species were found at both low and high elevation sites, while the other species were restricted to low (*C. parvicauda, C. zanzibari, C. losolos sp. nov., C. mwetiwel sp. nov.,* and *C. nansapw sp. nov.*) or high elevations (*C. ileile*). Temperatures showed the reciprocal pattern.

Considering forest type, we found that *C. pwilidak*, despite being the most abundant elsewhere, was absent from Botanical Garden, our one urban locality (0 of 15 worm-positive samples, despite its presence in 101 of 351 worm-positive samples overall; Fisher’s Exact Test *p* = 0.029).

The androdioecious species *C. tropicalis* and *C. briggsae* were present in all forest types, but were scarce in the mid-elevation rainforest, which was dominated by *C. pwilidak*. Other species were found too few times for an analysis across all forest types, but a binary classification into disturbed (agroforest, disturbed forest, urban) vs intact (rainforest, cloudforest) shows that *C. mwetiwel, C. nansapw,* and *C. losolos* are disproportionately found in disturbed forest (anova *p* = 0.008, 0.021, and 0.003, respectively; note that this forest classification is highly confounded with elevation and temperature). Species diversity was greatest in agroforest; a collection of 36 samples at a single agroforest site in the municipality of U, elevation 180-200 m, yielded eight species.

*Caenorhabditis* was present on a broad range of substrates. At the highest level of substrate classification, we found *Caenorhabditis* on 41% of rotting flower samples and 38% of rotting fruit samples, vs. 13% and 15% of rotting plant stems and fungi. *Caenorhabditis* was common in rotting fruits and flowers of trees native or endemic to Pohnpei, such as pwuhr (*Fagraea berteroana*), kotop (*Clinostigma ponapensis*), and nihn (*Ficus tinctoria*). Additionally, agroforest crops, including breadfruit, taro, banana, and star fruit, had high rates of successful *Caenorhabditis* isolation (Figure 4A). All of the species collected multiple times occurred on multiple different substrate types, including 15 different plant species plus fungus and leaf litter in the case of *C. pwilidak*.

**Figure 4.**
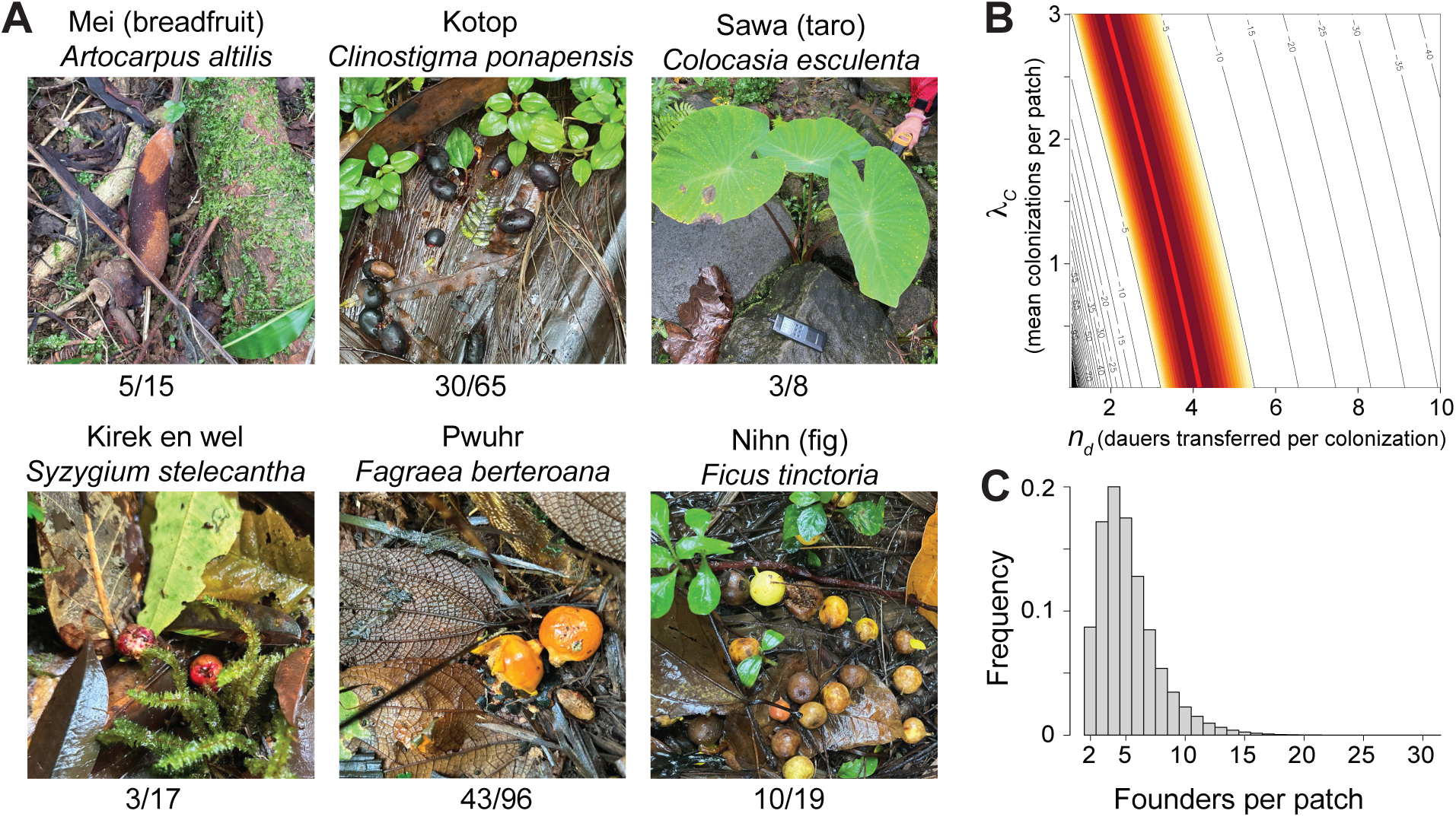
A. Common substrates for *Caenorhabditis* nematodes. The numbers below each photo report the number of *Caenorhabditis*-containing samples out of the total number of samples of that substrate type. **B.** Likelihood surface for the joint estimation of the number of colonization events per patch (*λ_C_*) and the number of founders transferred to a patch at each colonization event (*n_d_*) for *C. pwilidak*. The red line shows the ridge corresponding to the maximum log-likelihood (-1.87), with the colored region highlighting the 3-log 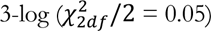 confidence interval. The ridge is well described by the quadratic 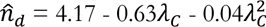 . **C.** The distribution of the number of founding worms per patch, for patches that successfully proliferate. Here we assume 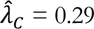, 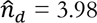, and sex ratio is 50:50. The mean of the distribution is 5.23.

We sampled 19 rotting substrates from elevated positions, including fruits and flowers still on their plants. Sixteen of these samples yielded worms, and four *Caenorhabditis*: in each case, exclusively *C. tropicalis* (n=1, 3, 5, and 5 individuals identified to species). The four samples were different substrate types – banana flower, ginger flower, iouiou fruit (*Canna indica*), and kirek-en-wel fruit (*Syzygium stelecantha*). For three of these samples, *C. pwilidak* was found in samples on the ground within a few meters. These data provide evidence that *C. tropicalis* may rely on different, potentially more volant, agents of dispersal than the other species of *Caenorhabditis* on Pohnpei.

### Properties of the patch colonization process

The data provide a method for inferring patch colonization parameters for *C. pwilidak*, our most common species. Seven of 101 samples that contained *C. pwilidak* included only a single worm of that species. These are patches that experienced only a single colonization event, and that event transferred only one worm. If we assume that colonization events are independent, and that the number of dauers transferred at each colonization event is Poisson distributed, we can use the data to jointly model colonization rate, with mean *λ_C_*, and dauer number per colonization, *n_d_* (File S1).

With only the observations of seven 1-worm patches out of 101 colonized patches, the two parameters cannot be estimated independently: the maximum likelihood forms a ridge (Figure 4B). The estimated number of dauers per colonization is 4.17 if colonizations are very rare, 3.50 if the mean number of colonizations per patch is 1 and 2.74 if the mean number of colonizations is 2. The mean number of colonizations per patch cannot be much greater than 2, given the prevalence of uncolonized patches. More specifically, from the observation that we sampled 400 patches to find the 101 occupied by *C. pwilidak*, we can independently estimate the colonization rate, under the assumption that the 400 patches are of comparable quality and accessibility and that colonizations are independent. Those numbers give an estimate of a mean of 0.29 colonizations per patch (95% CI 0.24 - 0.35). Conditioning on this estimate of *λ_C_*, we estimate 3.98 *C. pwilidak* dauer founders are transferred at each colonization event on average (CI = 3.20-5.00).

Because single-founder patches do not proliferate, and neither do patches with founders of only one sex, the mean number of founders of patches that contribute to subsequent generations is greater than the mean 3.98 dauers transferred per colonization. By simulation we estimate that proliferative patches have 5.23 founders on average (4.47 - 6.23 at the high and low end of the CI for *n_d_*), including cases where founders are contributed by multiple colonization events, and assuming an equal sex ratio (Figure 4C). Biased sex ratios (Huang *et al*. 2023) shift these values only slightly; for example, with an 80:20 bias, the average is 5.51 (4.69 - 6.58). Though these numbers convey unwarranted precision, the basic result that most colonizations involve multiple worms, but not many, is robust.

### Absence of C. elegans and C. inopinata

Although we collected 133 nematode-containing substrates from elevations above 500 m, 27 of them with temperatures below 22.5°C, we did not recover any *C. elegans*. Further, we found *C. tropicalis* in three of these 27 high, cool samples. Dense sampling in Hawaii suggests that *C. elegans* and *C. tropicalis* do not co-occur and have different habitat requirements, with *C. elegans* found in higher, cooler, less rainy locales (Crombie *et al*. 2019; Crombie *et al*. 2022a). The two species are isolated by latitude or elevation in other parts of the world as well. The presence of *C. tropicalis* in our highest, coolest samples suggests that Pohnpei currently lacks suitable habitat for *C. elegans*.

*C. inopinata*, the sister species of *C. elegans*, has a distinctive life history, living and reproducing within fresh figs of *Ficus septica*, transported from fig to fig by fig-pollinating *Ceratosolen* wasps (Woodruff and Phillips 2018). As *F. septica* does not occur on Pohnpei (Herrera *et al*. 2010), we searched for *Caenorhabditis* in fresh figs of nihn (*F. tinctoria*), the common local species. Dissected figs, verified to have undergone pollination, did not yield any *Caenorhabditis* worms.

### Fine-scale community composition

To complement our island-wide survey and to better understand the distribution of *Caenorhabditis* at finer scales, we densely sampled a single substrate type, rotting nihn figs from the forest floor beneath a single tree on Sokehs Ridge, elevation 246 meters. From each of three square-meter quadrats, we sampled a single fig from each of the 25 20-cm x 20-cm sectors (Figure 5A). These figs were selected to be at a matched, intermediate state of decomposition, based on color. We also sampled leaf litter from three of the sectors per quadrat (Figure 5B), and from the central sector of each, we sampled nine additional figs selected to span the range of decomposition states (Figure 5C). From each sample’s funnel yield, we estimated the total nematode density, regardless of clade, and then identified up to 10 *Caenorhabditis* individuals to species, or as many as were present if fewer than 10 (Figure 5).

**Figure 5.**
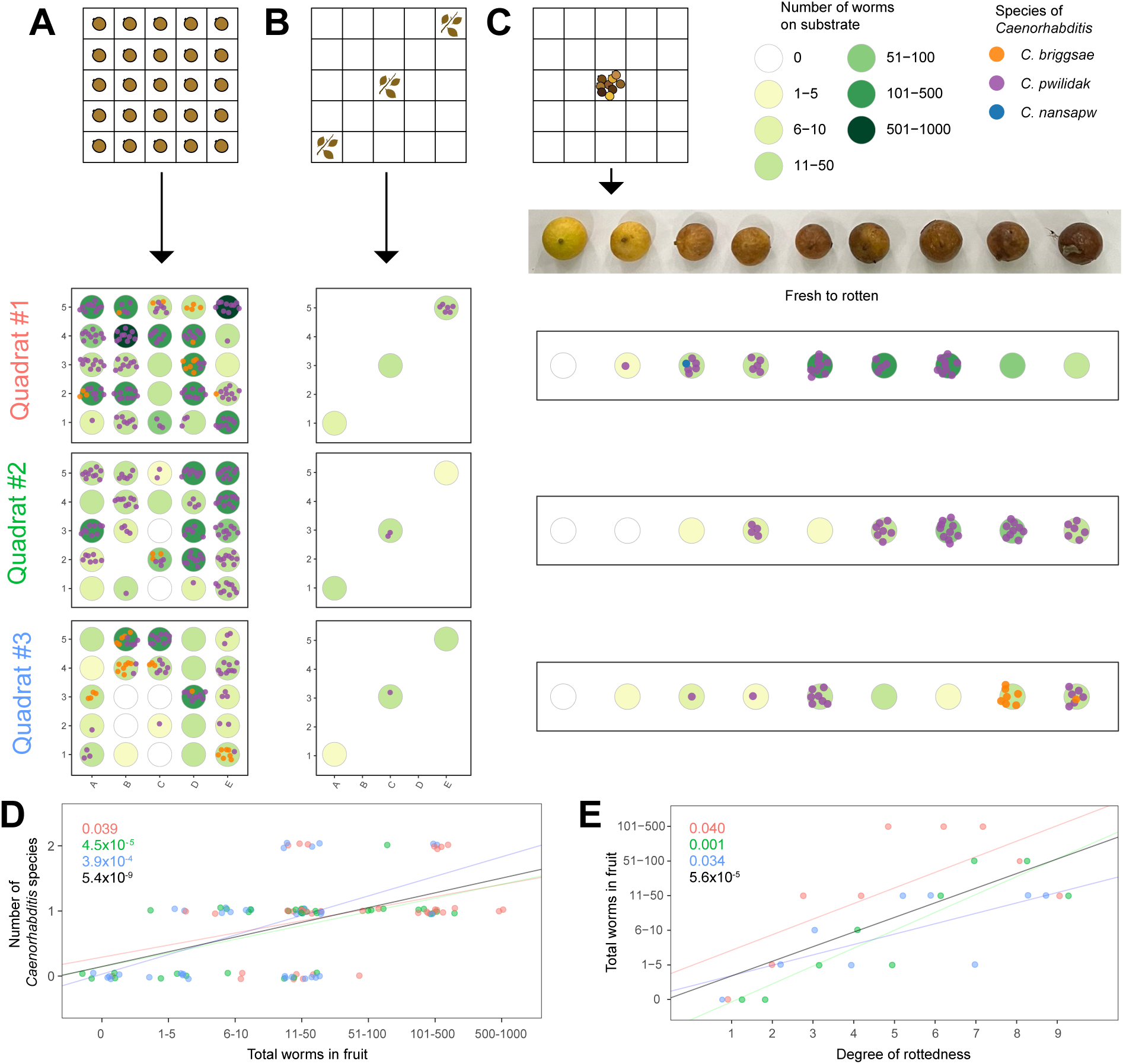
Fine-scale patterning of worm communities. In each of three replicate quadrats, we sampled (**A**) 25 similarly-rotten figs (color 5) from a grid of 25 squares, (**B**) leaf litter under three of the squares, and (**C**) nine figs across a range of rottenness from the central square of the grid. We found *C. briggsae* and *C. pwilidak* in many samples, and *C. nansapw* in one. Each dot represents an isofemale line or individual. One sample from quadrat 2 was lost. **D.** Figs with higher worm densities had more species of *Caenorhabditis*. The numbers report p-values for each quadrat individually and for the entire sample. Points are plotted with jitter in both axes. **E.** Worm densities were higher in figs that were further along the yellow-to-black spectrum. The numbers report *p*-values for each quadrat individually and for the entire sample.

Across these 111 samples (75 spatially arrayed figs, 27 color-range figs, and 9 leaf-litter samples), 75 contained *Caenorhabditis* (Table S6). From these we identified 482 *Caenorhabditis* isolates to species: *C. pwilidak* (n=425), *C. briggsae* (n=56), and *C. nansapw* (n=1). We hypothesized that we might see evidence for a priority effect, where each nematode-dense sample would be heavily dominated by a single species (De Meester *et al*. 2016; Ferrari *et al*. 2017). However, the proportion of worms in each sample that represent the most common *Caenorhabditis* species in that sample was not correlated with overall nematode density (*p* = 0.1). The total density of nematodes in a sample explained variation in the number of *Caenorhabditis* species present (*p* < 10^-8^): higher density meant more species (Fig 5D). Finally, presence of *C. pwilidak* and of *C. briggsae* is independent across the spatially sampled figs (Fisher exact test, *p* = 0.72), showing no evidence for a priority effect or interspecific aggregation.

Among the superficially identical figs in the spatially arrayed sample, 70% contained *C. pwilidak* and 19% *C. briggsae*. Assuming colonization events are independent, each fig received on average 1.2 colonizations (confidence interval 0.9-1.6) by *C. pwilidak* and 0.2 (0.1-0.3) by *C. briggsae*. These numbers imply that approximately 49% of figs occupied by *C. pwilidak* received multiple colonizations by that species, while the multiple-colonization rate for *C. briggsae* is 10%.

Fallen figs range in color from yellow (fresh) to black (Fig 5C). We observed a significant association between darkness and total nematode density (*p* < 10^-4^; Fig 5E), but not with the dominance of the most common *Caenorhabditis* species (*p* = 0.93), our test for priority effects, nor with the number of *Caenorhabditis* species (*p* =0.15).

To test the scale of spatial autocorrelation of worm populations among the color-matched figs (Figure 5A), we calculated Geary’s *c* (Geary 1954) on three different attributes – total number of worms, presence or absence of *C. pwilidak*, and presence or absence of *C. briggsae*. Geary’s *c* reports whether the spatial distribution of these attributes is clustered (<1), random (∼1), or uniformly dispersed (>1). Geary’s *c* for total number of worms on each fig was 0.87, 4.5 standard deviations below the mean of 100 permutations (*p* <3×10^-6^). This result is driven by differences between quadrats; when only comparisons within quadrats were allowed the pattern is not statistically significant (*p* = 0.51). We were similarly unable to detect any non-random distribution for the presence of the two species recovered, whether testing across quadrats (*C. pwilidak, p* = 0.82; *C. briggsae, p* = 0.69) or strictly within (*C. pwilidak, p* = 0.31; *C. briggsae p* = 0.39).

Sokehs Ridge is disturbed forest, and our island-wide survey also included 43 worm-containing samples there. These contained *C. briggsae, C. tropicalis, C. nansapw,* and *C. pwilidak*. The island-wide survey also included 19 samples of rotting nihn figs from five other regions; these yielded *C. briggsae, C. losolos,* and *C. pwilidak*. Including both the island-wide survey and the spatial samples, we recovered 5 different species of *Caenorhabditis* from rotting nihn, but curiously, never *C. tropicalis*, the second most abundant species in the survey. Of all *Caenorhabditis*-containing samples collected at Sokehs Ridge, *C. tropicalis* was found in 7/18 non-nihn samples and 0/79 nihn, consistent with a *C. tropicalis* substrate usage bias (Fisher Exact Test, *p* = 2×10^-6^).

### Phylogeny and biogeography

We assembled transcriptomes for each of the five new species and used the gene trees inferred from 2,955 proteins to generate a phylogeny incorporating 70 *Caenorhabditis* species (Figure 6; File S2).

**Figure 6.**
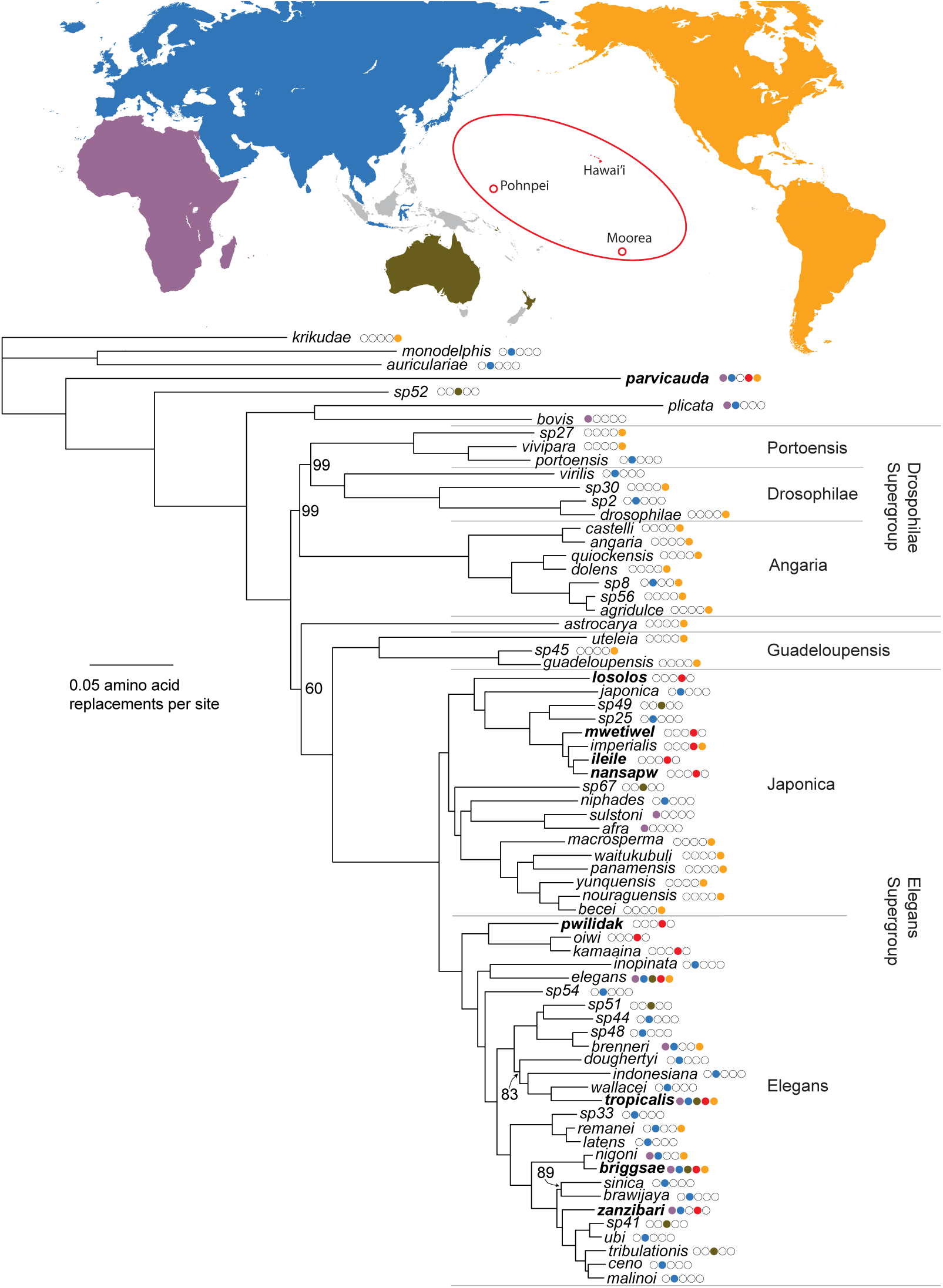
*Caenorhabditis* phylogeny based on 2,955 genes. Gene trees were estimated by maximum likelihood from protein sequences and the species tree estimated from the ensemble of gene trees. The species tree is rooted with the three species of the Auriculariae Group (Slos *et al*. 2017; Sloat *et al*. 2022). Support for each node, based on bootstrapping gene trees, is 100% except at five nodes, where the percentage of support is indicated in the figure. The nine species found in Pohnpei are shown in bold. The known distributions of each species are indicated by the circles to the right of the species names, using the five geographic regions designated in the map (see Methods for details). Gray regions on the map highlight islands of the southwest Pacific from which no data are available. Named species groups and supergroups are indicated.

The phylogeny differs from recent 51- and 60-species phylogenies (Fusca *et al*. 2025; Salome-Correa *et al*. 2025) only in the relative positions of *C. astrocarya* and Drosophilae Supergroup lineages as successive outgroups to the clade encompassing the Guadeloupensis Group and Elegans Supergroup.

Eight of the nine species observed on Pohnpei are within the Elegans Supergroup, scattered across many branches. The ninth species, *C. parvicauda*, is distantly related.

The five new species fall in two parts of the phylogeny. The most widespread and abundant species, *C. pwilidak,* is sister to the Hawaiian-endemic clade of *C. kamaaina* and *C. oiwi,* forming a Remote Oceania clade sister to the rest of the Elegans Group. The other four species fall within a small section of the Japonica Group, along with four known species: *C. japonica* (endemic to southern Japan: Kiontke *et al*. 2002; Yoshiga *et al*. 2013; Yoshiga 2018), *C. sp. 49* (Solomon Islands: evolution.wormbase.org), *C. sp. 25* (Singapore: evolution.wormbase.org), and *C. imperialis* (Moorea, French Polynesia, as well as Guadeloupe and Martinique, in the Lesser Antilles: Félix *et al*. 2014). *C. losolos* is the sister to the seven other species in this clade, while *C. mwetiwel*, *C. nansapw*, *C. ileile*, and *C. imperialis* are closely related to one another, with *C. nansapw* and *C. ileile* sister species.

*C. ileile,* collected at 671 m elevation, is unique in its apparent restriction to the Pohnpeian cloud forest. Given its close phylogenetic relationship to *C. mwetiwel, C. nansapw,* and *C. imperialis*, lowland species (<300 m) from Pohnpei and French Polynesia, we tested whether these species show differences in thermal tolerance. At 30°C, cultures of the three lowland species founded by a single pair of L4 juvenile male and female worms proliferated and exhausted their food supplies in the F_2_ generation, within 5 days (6/6 cultures per species). *C. ileile* cultures resulted in small F_1_ broods and no F_2_s, with resources remaining unconsumed after two weeks (6/6). Conversely, at 16°C, *C. ileile* and *C. imperialis* proliferated and exhausted their resources within two weeks, while *C. mwetiwel* and *C. nansapw* produced only F_1_s and no F_2_s (6/6 cultures per species in all cases).

The four previously named species found on Pohnpei are all known from multiple widely dispersed localities, around the globe (*C. briggsae, C. tropicalis, C. parvicauda*: Cutter *et al*. 2006; Gimond *et al*. 2013; Thomas *et al*. 2015; Stevens *et al*. 2019; Noble *et al*. 2021; Crombie *et al*. 2024) or across the Paleotropics (*C. zanzibari:* Stevens *et al*. 2019; Huang *et al*. 2023).

We inferred ancestral geographic ranges by maximum likelihood, and models with founder-event speciation are strongly favored over models that lacks such a scenario (Table S7, *p* < 10^-12^). These best-fitting models accommodate speciation events associated with long-distance dispersal, where the descendant species inherits none of its ancestral range (Matzke 2014). Because each species in the analysis occurs only in a single region, this dispersal-speciation parameter accounts for nearly all of the range shifts across the phylogeny, and alternative treatment of the other model parameters has little or no effect. Simultaneous dispersal and speciation is modeled as an event associated with branching events in the phylogeny, and consequently branch lengths are effectively irrelevant to the ancestral area reconstruction in this case.

Using the parameterization with the best fit (DEC+J; Table S7), the marginal ancestral-range probabilities at the nodes reveal uncertainty about the location of ancestors deep within the Elegans Supergroup (Figure 7). Remote Oceania receives the highest probability for the ancestral range of the last common ancestors of the Elegans Supergroup, the Elegans Group, and the Japonica Group (0.43, 0.61, and 0.40), but Eurasia is also plausible in each case (0.31, 0.37, and 0.32). The most recent common ancestor of *C. elegans* and *C. inopinata* is also plausibly Eurasia (0.66) or Remote Oceania (0.33).

**Figure 7.**
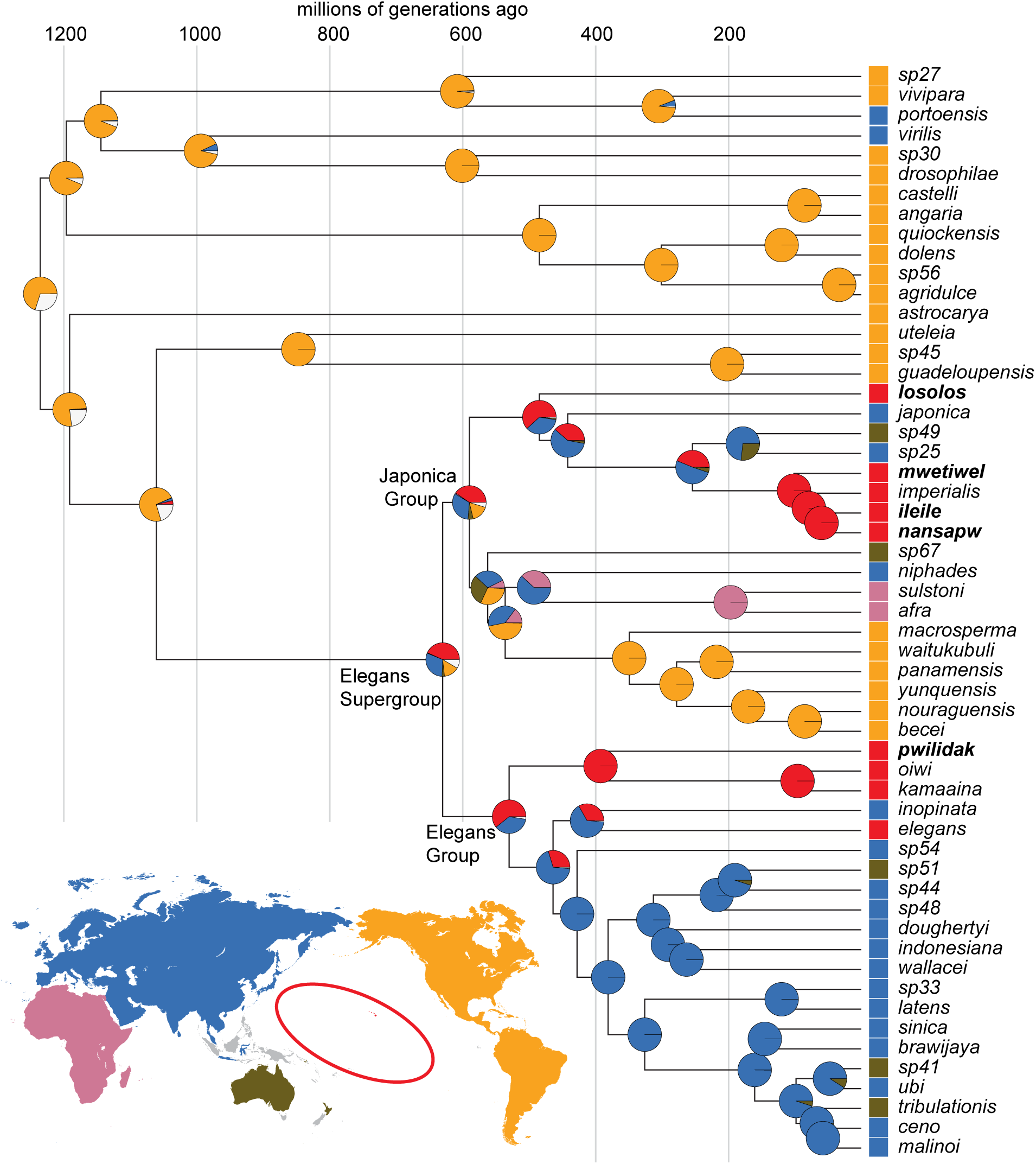
Maximum likelihood marginal reconstruction of ancestral species ranges, under the DEC+J model (see Methods for additional details). Each pie-chart records the marginal probability that the ancestor lived exclusively in one of the five areas, or, for pie regions that are white, in multiple regions. The lineage leading to the Elegans Supergroup likely emerged from the Neotropics, and the crown-clade diversification occurred later from an ancestor that may have lived in Eurasia or Remote Oceania. The timescale is adapted from Fusca *et al*. (2025).

## DISCUSSION

Our understanding of *C. elegans* at the cellular and molecular level has no match among animals, but our ignorance of its biogeography and natural history is an obstacle to the transformation of this species and its relatives into model powerful models for evolutionary and ecological questions. We intensively characterized the *Caenorhabditis* nematodes on a remote island, one uniquely suited to resolve outstanding questions about these worms, to lay foundations for ecological, biogeographic and evolutionary studies.

### The *Caenorhabditis* fauna of Pohnpei in context

We found nine species of *Caenorhabditis* nematode on the island of Pohnpei. Only two tropical forest sites have received comparable research attention, Barro Colorado Island in Panamá and Nouragues Natural Reserve in French Guiana, where six and nine species have been found, respectively (Ferrari *et al*. 2017; Sloat *et al*. 2022). Other well studied regions include the Hawaiian Islands, with five to seven species (two detected only by DNA sequence; Crombie *et al*. 2022a), and northern France, with five species (Félix and Duveau 2012; Félix *et al*. 2014). The high species diversity on Pohnpei is consistent with the latitudinal diversity gradient in *Caenorhabditis*, and with the global presence of *Caenorhabditis* in humid tropical forests revealed by opportunistic sampling around the world.

One surprise is that Pohnpei’s diversity is comparable to that found in continental tropical forest sites; this remote island’s *Caenorhabditis* community does not bear the signatures of distance from source populations that classical island biogeography predicts. This lack of island signature is also born out by the abundance of obligately-outcrossing species. Self-fertile organisms are expected to have an advantage in colonizing remote islands, as a single individual is sufficient to start a population (Baker 1955), and Pohnpei is a model island for testing this idea, known as Baker’s Law (Yomai and Williams 2021). We found that *C. briggsae* and *C. tropicalis,* the two species of self-fertile *Caenorhabditis* found in hot tropical regions, were present on Pohnpei, as they are in tropical forests around the world. The obligate outcrossing species *C. parvicauda* and *C. zanzibari* are also geographically widespread and may share attributes that predispose them to successful colonization (for example, low levels of inbreeding depression, or longer duration of viability in the dispersal-associated dauer state). Ancestors of all seven obligately-outcrossing Pohnpeian species must have arrived on Pohnpei as obligate outcrossers, given our understanding of the evolutionary origins of selfing from gonochorism in *Caenorhabditis* (Haag *et al*. 2018). A scenario that may be common in island-colonizing plants, where they arrive as selfers and subsequently evolve higher outcrossing rates (Baker 1967), is not applicable to these worms.

Overall, the paucity of selfing species in *Caenorhabditis* limits its power to test Baker’s Law, though an absence of *C. briggsae* or *C. tropicalis* would have been evidence against it. Hawaii, the other well studied remote island system, perhaps provides a better model. Hawaii is home to all three known selfing *Caenorhabditis*, and only four obligate outcrossers. Two of the outcrossers are sister species, and so the total number of colonization events implied by its fauna is just three each for the selfers and outcrossers. In this context, it seems that Pohnpei is not remote in the same way Hawaii is.

Like Hawaii, Pohnpei is home to multiple distantly related *Caenorhabditis* species, implicating independent colonization events, and also to sibling species, consistent with *in situ* speciation. One curious observation is that neither of the well studied continental rainforest sites, Barro Colorado and Nouragues, is home a to a pair of sister species. Pohnpei’s sister species *C. nansapw* and *C. ileile* provide a particularly attractive model for studies of speciation. These species are geographically separated into low-elevation and high-elevation domains, though *C. ileile* was only observed once, and they have different thermal tolerances that match their habitats. Speciation along elevational gradients is an expected pattern in remote islands (Gillespie *et al*. 2012). *C. ileile* is also our only cloudforest-endemic species. Pohnpei has only a few square kilometers of high-elevation cloud forest, and climate change renders this tiny region precarious. But even without climate change, the high elevations of Pohnpei have been gradually shrinking as the island erodes and subsides, on its geological route to atoll status. Oceanic islands give birth astride of a grave.

### Factors maintaining species diversity

Eight of the nine species on Pohnpei occur in the hot humid lowlands, and all eight were collected from a single agroforest site in U. The co-occurrence of so many species from a single guild, bacterivores that proliferate on rotting fruit and flowers, with little or no morphological differentiation, raises questions about the mechanisms that permit coexistence. Our data allow us to assess several plausible models, with a particular eye to the distinctive population biology of *Caenorhabditis* worms: their boom-and-bust dynamics that follow colonization of ephemeral resources patches (Félix *et al*. 2013; Ferrari *et al*. 2017; Sloat *et al*. 2022). This pattern implies the fitness of species and genotypes depends on a combination of within- and among-patch processes, with colonization rates playing a key role.

We used the frequency of patches containing single *C. pwilidak* worms to infer the average number of colonists per colonization: 3.98. This is consistent with several analogous estimates for other species, each using different methods. *C. elegans* genotype distributions in an orchard in France yielded an estimate of 3-10 founders per patch (Richaud *et al*. 2018). *C. inopinata* travels to new patches on fig wasps; direct counts found that each patch receives an average of 2.8 wasps, each carrying 0-6 worms (Woodruff and Phillips 2018). All of these data imply that patches start with very small numbers of individuals, and both biparental inbreeding and group selection are likely common features of *Caenorhabditis* biology. At the same time, most colonization events involve more than one worm, providing some explanation for *Caenorhabditis’* apparent escape from Baker’s Law.

All things being equal, species that are better resource competitors will drive poorer competitors to extinction in a homogeneous environment. In a patchy environment, a poor competitor can persist if it is better at colonizing new patches, gaining a head-start on proliferation before other species arrive to supplant it (Levins and Culver 1971; Tilman 1994). Colonization in *Caenorhabditis* involves phoresy, with larval nematodes in the dispersal-specialized dauer state hitching rides on passing invertebrates, and species differences in phoresy modes could contribute to differences in colonization rates. We observed that *C. tropicalis* was uniquely present in rotting fruits and flowers still on the plant, at positions elevated from the forest floor. It was also absent from rotting figs, where it should have been observed if it is distributed randomly among substrates. Both observations suggest that *C. tropicalis* may rely on different vectors to get to new patches, and its vectors may give it a colonization advantage for some fruits and flowers.

Absent differences in colonization ability or resource partitioning, species can coexist if patchy environments result in patchy species distributions. If individuals of each species are aggregated, with patch founders sharing patches primarily with founders of the same species, the increase in intraspecific competition and decrease in interspecific competition, relative to the continuous-landscape case, creates conditions for coexistence (Shorrocks 1990; Hartley and Shorrocks 2002; Czekanski-Moir and Rundell 2019).

The intraspecific aggregation model of coexistence has been successful in describing empirical patterns in multispecies guilds of arthropods on ephemeral resource patches (Ives 1991), and it appears to be compatible with our data from the spatial arrays of superficially identical figs in quadrats at Sokehs Ridge. We can quantitatively (if imprecisely) assess conformity with the requirements of the intraspecific aggregation model by estimating the intraspecific aggregation index, *J*, of Ives (1991). Assume that each patch receives a small, Poisson-distributed number of colonization events; we estimated 1.2 for *C. pwilidak* and 0.2 for *C. briggsae* in Sokehs Ridge figs.

Assume that each colonization delivers a small, Poisson-distributed number of founders for each species; we estimated 3.98 for C*. pwilidak*. Using these numbers in a million simulated Sokehs Ridge experiments (File S1), *J_pwilidak_* averaged 0.80 (standard deviation = 0.19), an 80% increase in intraspecific aggregation over independent assignment of individual founders to figs. If we assume that the 3.98 founders per colonization also applies to *C. briggsae* at Sokehs Ridge, the estimate of *J_briggsae_* is 4.84 (sd = 1.79). Finally, assume that the colonization process is independent for each species, which is supported by our observation that *C. pwilidak* and *C. briggsae* were distributed independently among the figs. Applying Ives’ (1991) equation 11, we estimate that intraspecific aggregation at Sokehs Ridge reduces the interspecific competition between *C. briggsae* and *C. pwilidak* by 89% (sd = 19%). Depending on the magnitudes of competition coefficients, this level of aggregation could facilitate long-term stability of a community that would otherwise not persist.

Though the specific numbers surely require a grain of salt, the general result is that multiple dauers per colonization plus independent colonizations results in a population structure that comports with the requirements of the aggregation model of coexistence.

In the extreme of the aggregation model, competitors can find themselves in their own patches. Many figs at Sokehs Ridge contained only one species or the other, and some figs contained neither, providing empty habitat patches (Figure 5A). Eight percent of the figs contained no nematodes, and 26% contained no *Caenorhabditis*. We also found no spatial autocorrelation in worm density or species presence across the 25 sections within each of our 1-m^2^ quadrats, consistent with dispersal distances of greater than tens of cm and a lottery-like patch colonization process rather than a simple moving front.

Finally, we note that other mechanisms of competition or resource partitioning may be important for Pohnpei’s *Caenorhabditis* community, given the frequent co-occurrence of species within individual rotting substrates (Fig 3A, Fig 5A). An emerging mechanism for persistence of species sharing the same resources is reproductive interference, which can allow weaker resource consumers to maintain their numbers by damaging population growth in their competitors; this phenomenon has been experimentally demonstrated in *Caenorhabditis* nematodes (Ting and Cutter 2018; Schalkowski *et al*. 2024). We found that the cooccurring *Caenorhabditis* species of Pohnpei readily mate with one another under no-mate-choice experimental conditions, and consequently reproductive interference may play a role in their population biology.

Species coexistence may also depend on resource partitioning that we failed to observe. Co-occurring species showed some level of habitat differentiation, with *C. pwilidak* dominating mid-elevation rain forest, and other species – *C. losolos, C. mwetiwel, C. nansapw,* and *C. zanzibari* – occurring only in disturbed habitats: agroforest in Kitti and U, the botanical garden in Nett, and adjacent to a dirt service road in Sokehs. The coincidence of eight species at one site in U may therefore not represent an equilibrium coexistence state but a sink for some species that will be outcompeted locally but persist in resource refuges elsewhere.

Our data from Pohnpei are purely observational, and experimental approaches to the study of patch colonization hold promise for resolving these questions (Ferrari *et al*. 2017; Sloat *et al*. 2022).

### Pohnpei in global *Caenorhabditis* biogeography

*Caenorhabditis* biogeography is in its infancy. The very first glimpses of geographic patterns emerged only in 2011, when fewer than 40 species were known (Kiontke *et al*. 2011). The nine species from Pohnpei, placed into the context of *Caenorhabditis* phylogeny and geography, help us frame and address two major questions about the history of this clade. We first ask where the ancestors of *C. elegans* and the Elegans Supergroup species lived and when. Then we ask how the nematodes got to Pohnpei, considering both the biology of the worms and the history of the Pacific. Finally, we use the biogeographic data to make predictions about the worms in regions yet to be sampled.

The data support a history in which the Elegans Supergroup, with its diverse species spread across the world, arose from an American emigrant that moved westward, into and across the Pacific.

American origins for terrestrial species in Oceania and the western Pacific are not unprecedented, despite the vast distances that separate those regions. Analyses of the floras of Hawaii and Micronesia suggest that both received substantial input from New-World sources (Price and Wagner 2018; Demeulenaere and Ickert-Bond 2022). Animals provide parallel examples, as in the case of Zalmoxid opiliones, arachnids inferred to have crossed from the Neotropics to the west pacific in the Cretaceous (Sharma and Giribet 2012). More recent cases include Hawaiian insects, which hail largely from the Americas (Gillespie *et al*. 2012), and the Fijian iguana, whose ancestors rafted across the Pacific (Scarpetta *et al*. 2025).

The data are equivocal about the site of diversification of the Elegans Supergroup. One plausible model places the most recent common ancestor of the supergroup at the western edge of the Pacific, with multiple subsequent colonizations of Remote Oceania. Under this model, *C. elegans* are in Hawaii because their Asian ancestors dispersed eastward across the Pacific. An alternative model, one with slightly stronger support from our data, suggests that the Elegans Supergroup ancestor lived and diversified within Remote Oceania, giving rise to Pohnpeian and Hawaiian lineages, including *C. elegans*, and ultimately colonizing of East Asia and Australia multiple times. Oceania is then a source for the Old World, an evolutionary layover for American worms heading west.

The hypothesis that the American emigrant ancestor of the Elegans Supergroup diversified in Oceania rather than Asia is suggested by the ancient clade of Pohnpeian and Hawaiian worms – *C. pwilidak, C. kamaaina,* and *C. oiwi* – and by the other deeply rooting Oceanian species in both the Japonica and Elegans groups – *C. losolos* and *C. elegans* respectively. These observations are consistent with a stepping-stone history of migration (Price and Wagner 2018). Examples of Oceanian species with sources in distant islands rather than continents are accumulating, such as the sandalwood and *Melicope* trees and shrubs in Micronesia and eastern Polynesia that have Hawaii as their source (Harbaugh and Baldwin 2007; Harbaugh *et al*. 2009), or the Tahitian land snails whose closest relatives are all from Hawaii’s Big Island (Rundell *et al*. 2004). More generally, the traditional view of remote islands as species sinks is yielding to emerging phylogenetic data that islands contribute as sources as well (Hembry *et al*. 2021; Keppel *et al*. 2024).

Notwithstanding these possibilities, the relative proximity of New Guinea and the Philippines – regions for which we currently lack data – provides a proven nearby source for Micronesian taxa (Demeulenaere and Ickert-Bond 2022; Keppel *et al*. 2024). In a metaanalysis of the biotas of Pacific islands, Keppel *et al*. (2024) found a strong relationship between proximity to the source area and the number of colonization events – nearby areas are the usual sources. Their underlying data represent a wide range of taxa, primarily plants, with modes of dispersal that may render them poor predictors for *Caenorhabditis* nematodes. Further, data for Pohnpei are sparse, and generalizations about Micronesia may not apply.

The timescale for *Caenorhabditis* evolution is poorly constrained, due both to the absence of fossil calibration data and the exceptional high rate of molecular evolution in the clade (Cutter 2008); comparisons of sequences from even close relatives reveal substitutional saturation. Recent work has heroically circumvented these problems and estimated the ages of nodes in the *Caenorhabditis* phylogeny in terms of numbers of generations (Fusca *et al*. 2025), which can then be converted to dates with information about generation times. Unfortunately, the number of generations per year is unknown for *Caenorhabditis* worms, and it likely varies enormously among species and populations (Cutter 2008). *Caenorhabditis* worms can persist in the dauer state for months, when conditions are poor, generating a sort of seed bank (Barriere and Felix 2005; Cutter 2015), and under good conditions generation time varies as a function of species, diet, and temperature. Fewer than 10 and more than 100 generations per year are both plausible.

Applying the timescale from Fusca *et al*. (2025) to our phylogeny (Figure 7), the most recent common ancestor of the Elegans Supergroup lived approximately 640 million generations ago, while the putatively American common ancestor of the Elegans Supergroup and Guadeloupensis Group lived approximately 1080 million generations ago; these numbers correspond to dates ranging from 64 and 108 million years at ten generations per year to 6.4 and 10.8 million years with 100 generations per year. Reality likely falls between these extremes.

The locations of islands and continents in western Pacific during the course of the Cenozoic are somewhat uncertain, as this region has been the site of exceptional tectonic dynamism; an area equivalent to one seventh of the entire surface of the earth has subducted there during the last 50 million years (Wu *et al*. 2016). Pohnpei formed approximately 8 million years ago, too late to have been the site of colonization directly from America under the long-generation-time end of the inferred evolutionary timescale but just plausible under the short-generation-time estimates.

However, worms could have migrated westward one island at a time, using stepping stones that have long since sunk beneath the waves (Nunn 2008; Claridge *et al*. 2017; Hembry *et al*. 2021; Keppel *et al*. 2024). When Pohnpei formed, it was further from Asia and Australia than it is now, flanked distantly by the high island of Chuuk to the west and the atolls of the Marshall Islands to the north and east. Though the current major Hawaiian Islands were not yet emergent, the Hawaiian chain has likely had elevations above 500 m continuously for the last 30 million years (Price and Clague 2002), and the Marshalls date back to the Cretaceous—long enough ago to have provided safe harbor to westward migrant worms even under the long-generation-time scenario. The positions of Pohnpei, Chuuk, the Marshall Islands, and Hawaii have all been stable relative to one another, as they sit together on the Pacific plate, which is slowly drifting westward, bringing them closer to Asia and the western islands of Micronesia—Yap, Palau, and the Marianas.

One potential constraint on timing is the apparent *in situ* speciation between lowland *C. nansapw* and highland *C. ileile*. If this event happened on Pohnpei, as we infer, it must be less than 8 million years ago. The molecular clock places their ancestor approximately 60 million generations ago (or a bit less, accounting for the difference between the sequence coalescence time and the population divergence time). This corresponds to 6 or 0.6 million years at 10 or 100 generations per year, but effectively excludes the possibility of many fewer than 10 generations per year. If we suppose that the common ancestor of *C. mwetiwel, C. imperialis, C. nansapw,* and *C. ileile* also lived on Pohnpei, less than 8 million years ago, that clade’s age of ∼102 million generations would constrain the life cycle to more than 12 generations per year. These very weak constraints underscore the challenges of dating *Caenorhabditis* evolution.

One finding that is robust to the vagaries of the molecular-clock estimates is that the Elegans Supergroup emerged well after the opening of the Atlantic Ocean, supporting the inference that it reached Asia via long-distance dispersal westward from America across the Pacific, rather than via Gondwanan vicariance; the embedding of the Elegans Supergroup within many Neotropical outgroups further argues for a post-Gondwanan timescale.

Having spread across the Pacific and East Asia, the diversifying Elegans Supergroup then gave rise to three lineages that traveled further, arriving in Africa (the clade of *C. sulstoni* and *C. afra*), India (*C. doughertyi*), and back to the Neotropics (a clade of six species in the Japonica Group). Several other species in the Elegans Group spread across the globe, likely with help from people.

### How did *Caenorhabditis* get to Pohnpei?

Every species in Pohnpei descends from ancestors that dispersed across open ocean. The long-distance dispersal events that generate the faunas of oceanic islands involve just four mechanisms: drifting on the winds, rafting on the ocean, hitching rides on birds or bats, and traveling with humans (Gillespie *et al*. 2012; Demeulenaere and Ickert-Bond 2022). Any of these mechanisms could carry *Caenorhabditis* nematodes, which can withstand starvation and severe desiccation in the specialized dauer state (Erkut *et al*. 2011). Careful experimental data show that *C. elegans* dauers readily survive two months (Klass and Hirsh 1976), and up to four under ideal conditions (Kaptan *et al*. 2020), but the common experience of *Caenorhabditis* researchers is that cultures regularly recover from even longer periods of neglect. In one unplanned experiment, cultures of tropical *C. briggsae* and *C. tropicalis* had a high rate of survival after six months in parafilmed petri dishes (Sloat *et al*. 2022). To interpret Pohnpei’s fauna, and to make predictions about *Caenorhabditis* in regions not yet studied, we consider each of the four dispersal mechanisms, in the context of possible source geographies.

#### Arrival by air

*Caenorhabditis* worms routinely travel phoretically on insects and other terrestrial arthropods (Kiontke 1997; Sudhaus *et al*. 2011; Yoshiga *et al*. 2013; Frezal and Felix 2015; Woodruff and Phillips 2018; Sun *et al*. 2022), which can be blown to remote islands. Ship-based entomology has shown that Pacific skies are full of wind-blown insects (*e.g.*, Holzapfel *et al*.1978), and a large fraction of the Hawaiian insect fauna arrived from the Americas on the prevailing winds (Gillespie *et al*. 2012), which continue southwestward to Pohnpei (Bosserell *et al*. 2015). Occasionally Pohnpei also experiences winds that blow west to east, particularly during El Niño (Lander and Khosrowpanah 2004).

Cyclones from the east provide an air route for larger animals, including snails (Cowie and Holland 2006), which can also vector *Caenorhabditis*. Pohnpei is close enough to the equator that cyclones are rare, however (Lander and Khosrowpanah 2004), and there are few potential source islands on cyclone paths to Pohnpei.

Though insect vectors are the most plausible means of aerial arrival, many nematodes can travel solo as aerial plankton (Ptatscheck *et al*. 2018). Long-distance travel by this mode likely requires anhydrobiosis, common in many groups of nematodes but not in *Caenorhabditis* (but see Erkut *et al*. 2011).

Overall, the wind route predicts that Pohnpei’s *Caenorhabditis* will have their origins to the Northeast, and the immigrant species will have phoretic relationships with insects that show similar biogeographic histories.

#### Arrival by sea

Rafting is slower than flying, but it provides another potential route for *Caenorhabditis* nematodes (Gillespie *et al*. 2012). An empirical model of debris drifting in the ocean (van Sebille 2014), parameterized with data from more than 17,000 drifting buoys, shows that debris that reaches Pohnpei after two or four months at sea is most likely to have originated to the south, on the north coast of western New Guinea or in the nearest Caroline and southern Marshall Islands. Rafts at sea for longer – ten months or a year -- could easily arrive from the Philippines or the coast of South or Central America. The experimental data underlying these models largely capture the behavior of the uppermost tens of meters of water; a worm-bearing raft of debris would also be influenced by winds, which will generally favor sources to the east. In addition, though Pohnpei’s setting north of the equator has been stable for its whole history, the oceanographic regime is likely to have changed significantly over its 8-million-year lifetime.

Pohnpei’s worms can also be a source – a stepping stone – to sites further west. The drift model (van Sebille 2014) shows nonnegligible probabilities for debris from Pohnpei to reach Chuuk and Guam within four months, and Yap, Palau, and the Philippines within six. Wind is likely to accelerate westward rafting from Pohnpei. Currents do not provide a likely route from Pohnpei to Hawaii, nor from Hawaii to Pohnpei.

#### Birds and bats

Flying vertebrates can carry animals to remote islands under their own power, escaping the geophysical constraints of wind and water. Worms can travel in or on snails (Chen *et al*. 2006; Petersen *et al*. 2015; Sudhaus 2018), which themselves can be carried in or on birds (Wada *et al*. 2012). *Caenorhabditis* could also, perhaps, travel freely in bird or bat intestines; the best evidence is that four worms, likely from the Elegans supergroup based on male tail anatomy, were recovered alive from the intestines of a plumbeous water redstart in Taiwan in 1959 (Schmidt and Kuntz 1972). Many bird species are widely distributed across the Carolines and beyond, and migratory birds using flyways in Micronesia are traveling among Old-World sites (Gillespie *et al*. 2012; Demeulenaere and Ickert-Bond 2022). In general, passage on flying animals is expected to homogenize island faunas in the western Pacific. Pohnpei’s endemic flying vertebrates have largely Old-World origins, including the iconic endemic lorikeet (serehd) and flying fox (pwehk) (Almeida *et al*. 2014; Schweizer *et al*. 2015), but their endemism is evidence that these do not provide regular inter-island passenger service for worms.

Hawaii’s flora arrived largely in or on birds, and largely from North American flyways (Price and Wagner 2018). These flyways help explain biotic similarities between Hawaii and the Marquesas far to its south, but they do not provide a route between Hawaii and Pohnpei. If birds played a role in the *Caenorhabditis* link between Hawaii and Pohnpei, it likely involved an intermediate step.

#### People

The fourth mechanism of island colonization is travel with humans. Each of the four previously described species we found on Pohnpei is widely distributed across multiple continents, and travel with humans is a likely (though not exclusive) scenario. If these species have been introduced to Pohnpei by people, the possible dates of introduction are somewhat constrained.

Pohnpei has been inhabited for approximately two thousand years (Athens 2018), and for most of its history, it had little exchange with other islands (Hanlon 1988). American and European traders first arrived in the 1820s, followed by American missionaries in the 1850s. Intentional agricultural introductions began immediately. Later occupations by Spain, Germany, Japan, and the United States resulted in more intensive plant introductions, particularly during the German (1899-1914) and Japanese (1914-1945) occupations (Ragone *et al*. 2001). More than 430 new plants were introduced, imported from more than 30 regions spanning the globe, particularly Japan, Indonesia, New Guinea, Hawaii, and Taiwan, but also California, Europe, and elsewhere (Ragone *et al*. 2001).

The center of plant introductions was the Pohnpei Agricultural Station in Kolonia. We sampled from the site of the station, now the Pohnpei Botanical Garden, and we recovered three of the putative introduced species there: *C. briggsae, C. tropicalis,* and *C. parvicauda*. We also found new species *C. mwetiwel* and *C. nansapw* there, but not *C. losolos, C. ileile,* or *C. pwilidak*; the latter was so abundant everywhere else on the island that its absence from the Botanical Garden is statistically notable. All of the species found at the Botanical Garden were also found in other parts of the island. Forthcoming population genomic data for *C. briggsae* and *C. tropicalis* isolates may help narrow the geographic sources and ages of their hypothesized introductions.

### Where next?

This work was motivated in part by the goal of finding a close sister species for *C. elegans*. The closest species currently known is *C. inopinata*, a species from Taiwan and the Ryukus with a highly derived anatomy and life history (Woodruff and Phillips 2018). At the molecular level, genes of *C. elegans* and *C. inopinata* are about four times as diverged as those of human and mouse (Mouse Genome Sequencing *et al*. 2002; Kanzaki *et al*. 2018). Their most recent common ancestor lived an estimated 418 million generations ago; for comparison, the most recent common ancestor of humans and chimpanzees is inferred to have lived 0.25 million generations ago (assuming 6 million years and 25-year generations: Langergraber *et al*. 2012; Wang *et al*. 2023; Yoo *et al*. 2025).

We focused on Pohnpei because it is home to the most plausible *C. elegans* habitat in the region between Hawaii and Taiwan. Though we found neither *C. elegans* nor a new sister species, our findings reinforce the idea that the islands of the Pacific are a promising place to continue the search. Cool cloud forests occur on several other islands in remote Oceania – in the Cook Islands, French Polynesia, Samoa, and Fiji (Merlin and Juvik 1995). Kosrae, Pohnpei’s younger neighbor 500 km to the southeast, is not as high as Pohnpei but it has 0.7 km^2^ of cloud forest on its summits (Whitesell *et al*. 1986). Lower elevation sites might also hold the key, on the theory that *C. elegans’* restriction to cool conditions is derived, much as *C. ileile*’s appears to be; in that case, the Marshall Islands, Chuuk, Yap, Palau, and Guam are all older islands that could retain descendants of ancestors recently shared with *C. elegans*. Finally, of course, the massive expanses of Papua New Guinea, Indonesia, and the Philippines have enormous elevational ranges and diverse forests, and the absence of data from those regions limits the strength of our biogeographic inferences.

Hypotheses generated by this work will find their tests in the yet undiscovered *Caenorhabditis* faunas of these regions.

## DATA AVAILABILITY STATEMENT

Sample collection data are reported in Supplementary Tables. Type cultures of each new species are deposited at the *Caenorhabditis* Genetics Center, and cultures of all isolates are available from the authors. Ribosomal DNA sequences have been deposited in Genbank with identifiers PP955316-PP955322. RNAseq data and transcriptome assemblies are deposited with NCBI under BioProject ID PRJNA1128046.

## SUPPLEMENTARY MATERIAL

## Appendix Descriptions of five new species

### Supplementary Tables

**Table S1. Transcriptomic Data.** For each species, the table lists the BUSCO annotation of the transcriptome (percent complete, single copy, duplicate, fragmented, and missing), the identity of the sequence reference strain, the legacy species number for species that are or were previously known by temporary species numbers, the source of the sequence data (*e.g., Caenorhabditis* Genome Project CGP), the reference for the data, the nature of the data (transcriptome or genome sequence), and the file name for the source data.

**Table S2. Biogeographic Data.** For each species, the table lists the legacy species number, the species name, its known geographic distribution, and a citation for the distribution information. The remaining columns record 1 or 0 for the presence or absence of each species from the specified geographic region, as shown in Figures 6 and 7, and indicate whether the species is included in the biogeographic analysis. Species were excluded if they were globally distributed, if their distributions were ambiguous (*e.g., C. sp. 2*), or if they fall outside the clade defined by the Elegans and Drosophilae Supergroups.

**Table S3. Survey Results.** This table provides sample-level information for each of the substrates examined as part of our survey of the island. It includes the 400 rotting samples at the core of the survey, and an additional 14 non-rotting samples including fresh figs and live invertebrates. The columns list, for each sample, the sample ID (corresponding to the barcoded sample bag), the Collection Set (one for each locality-day), the C-plate summary (reporting at a high level what worms if any emerged from each sample’s Baermann Funnel extraction onto a petri dish, the C-plate), the latitude, longitude, and elevation of the sample, the number of *Caenorhabditis* worms identified to species level, whether the sample included gonochoristic *Caenorhabditis* worms of a single sex only (precluding establishment of a culture), and then a series of columns reporting presence (1) or absence (0) of each species of *Caenorhabditis*. The remaining columns report the general substrate class, the substrate identity, whether the sample was rotting material (1 for yes, 0 for no), whether the sample was found on the ground or at an elevated position, the Pohnpei-specific vegetation type, the general landscape classification (following *Caenorhabditis* conventions), the substrate temperature, the ambient temperature, and the collection date.

**Table S4. All *Caenorhabditis* Isolates.** This table provides worm-isolate-level information for each of the individuals isolated from each of the 400 survey samples and 111 fig and leaf-litter samples from the spatial sampling arrays at Sokehs Ridge. For each isolate, the table lists the Isolate ID (Sample ID as in Table S3 with the addition of an upper-case letter for the Isolate ID), the species of *Caenorhabditis*, the QG number that represents the unique strain name for the cryopreserved culture established from the isofemale or isohermaphrodite isolate (following *C. elegans* genetics convention, QG indicates the laboratory and strains are assigned numbers sequentially), the Sample ID, a Yes or No as to whether the sample is part of the Sokehs Ridge spatial sample set, and then the Collection Set, latitude, longitude, and elevation, as in Table S3. The remaining columns report the general substrate class, the substrate identity, whether the sample was rotting material (1 for yes, 0 for no), whether the sample was found on the ground or at an elevated position, the Pohnpei-specific vegetation type, and the general landscape classification.

**Table S5. Mating Tests.** This sheet contains four tables, each describing the results of experimental test crosses among closely related strains or species. The cross data allow us to establish *C. ileile, C. nansapw, C. mwetiwel,* and *C. pwilidak* as biological species incompatible with closely related species, and they show that Pohnpeian isolates QG4816 and QG5228 are compatible with known isolates of *C. zanzibari* and *C. parvicauda*, respectively.

**Table S6. Spatial Sampling.** This table provides sample-level information about the 111 figs and leaf-litter samples collected from the spatial arrays shown in Figure 5. One sample, the fig from quadrat 2, position B2, was lost and is not reported here. For each of the remaining samples, the table shows the Sample Name, the sample ID in C-plate format, the number of individuals identified as *C. pwilidak, C. briggsae,* or *C. nansapw*, and the total number of *Caenorhabditis* species found among the identified worms. The WormDensity column reports an ordered factor describing the overall number of worms recovered from the sample’s Baermann Funnel, as follows: 0, 0 worms; 1, 1-5 worms; 2: 6-10 worms; 3: 11-50 worms; 4: 51-100 worms; 5: 101-500 worms; 6: 501-1000 worms. The remaining columns report the sample’s quadrat (1, 2, or 3), row, column, and type (fig or leaf litter). The 9 figs sampled from the central position of each quadrat to represent a range of rottenness levels are designated by “R” as their value under “quad_column,” and their rottenness level is indicated by the values 1-9 in the under “quad_row.”

**Table S7. Biogeographic Analyses.** This sheet contains two tables describing statistical results from biogeographic models. The upper table lists the negative log likelihoods for each of six models, along with their parameter estimates. Models incorporating jump speciation (+J) confer much higher likelihoods than those without, and the DEC+J model confers the highest likelihood. The lower table tests the significance of incorporating the J parameter, in each case finding that it significantly improves the likelihood.

### Supplementary Files

**File S1. Statistics of patch colonization.** This is a .R file containing scripts that estimate parameters of patch colonization and simulate populations to estimate aggregation statistics.

**File S2. *Caenorhabditis* phylogeny.** This is a Newick tree file containing the unrooted species tree estimated in Astral from 2955 gene trees. The branch lengths were then estimated in iqtree using the concatenated amino-acid sequences of 1,913 single-copy orthologs. This is the tree plotted in figure 6.

**File S3. Ultrametric tree for biogeographic analysis.** This is the input Phylogeny file for BioGeoBEARS analysis. It is a Newick tree file containing the rooted ultrametric species tree, as plotted in figure 7.

**File S4. Biogeography file.** This is the input Geography file for BioGeoBEARS analysis. The underlying data are sourced and adapted as described in Table S2.

## ACKNOWLEDGEMENTS

We gratefully acknowledge the assistance of our guides and consultants in Pohnpei, Holden Pelep (Kitti municipality) and Zelnick Moses (U municipality). We are grateful for support and assistance from Roseo Marquez and Tamara Greenstone Alefaio (Micronesia Conservation Trust), Eugene Joseph (Conservation Society of Pohnpei), Hubert Yamada (Pohnpei State Resources and Development), Vanessa Fread and Dave Mathias (FSM Resources and Development), Eugene Eperiam (Division of Natural Resource Management), Chief Minister Rufino Primo (U Municipal Government), and Chief Mihkel Bernardo, Ryan Agrippa, and Kenny Bernardo (U). Thanks to Micronesian Productions for the translation of the summary into Pohnpeian. We thank Derin Çaglar for help in the lab, NYU GenCore, supported by the Zegar Family Foundation, for sequencing support, and NYU HPC for computational support. We thank the *Caenorhabditis* Genetics Center (supported by NIH P40 OD10440) and the Fitch Lab at NYU for worm strains, WormBase and the *Caenorhabditis* Genomes Project for data, and the researchers who contributed strains and information to the CGP and to the worm evolution community wiki. Thanks to Manpreet Katari, Ryan Baugh, Zoltan Szuts, Erik Andersen, Justin Bernstein, and Jesse Czekanski-Moir for helpful advice. Apologies to Samuel Beckett for borrowing “birth astride of a grave” for island biogeography. The laboratory and computational parts of this work were supported by NIH R35GM141906.

## Appendix

### Descriptions of five new species

Following the approach of Félix *et al*. (2014), and in recognition that many *Caenorhabditis* species are morphologically cryptic, we employ experimental crosses to diagnose species according to the Biological Species Concept. Photographs of male tails are provided in Figure A1 in accordance with the International Code of Zoological Nomenclature guidelines for species descriptions.

The electronic edition of this article conforms to the requirements of the amended International Code of Zoological Nomenclature and new names contained here are available under that Code from the electronic edition of this article. This published work and its nomenclatural acts are registered in ZooBank. The ZooBank LSIDs (Life Science Identifiers) can be resolved and the associated information viewed through any standard web browser by appending the LSID to the prefix ‘‘http://zoobank.org/’’. The LSID for this publication is: urn:lsid:zoobank.org:pub:F8CC909D-582B-4EAC-A46F-E326F00E6F01

### Methods

Worms were cultured at 25°C on NGMA plates seeded with OP50-1 *E. coli* bacteria. Adult males, 1 day post L4, were heat-killed and mounted on 4% agar pads on microscope slides and imaged on a Zeiss AxioImager M2 microscope with DIC optics and EC Plan-Neofluar 100x/1.3 objective.

**Table A1.**
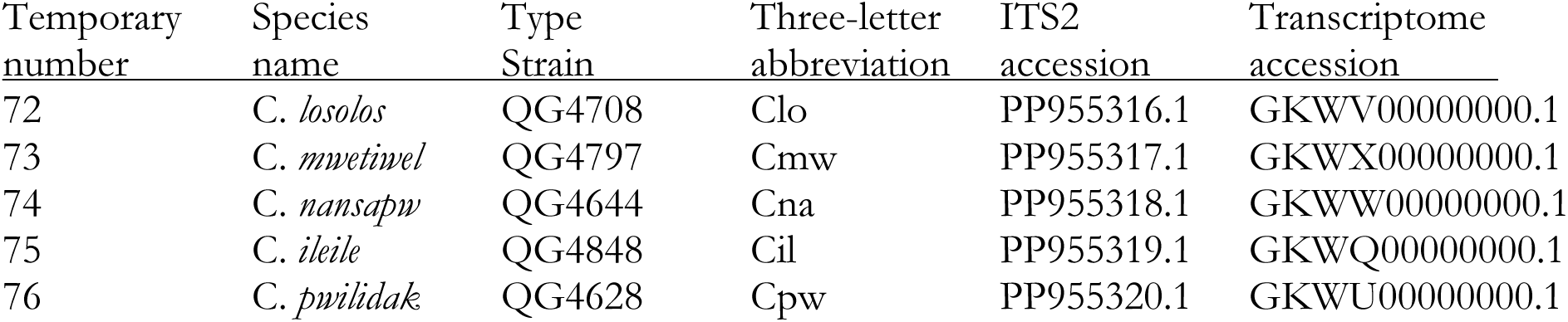
Summary of new species.

#### Species Declarations

***Caenorhabditis losolos* Rockman, Tintori, Nguyen, & Yomai *sp. nov*.**

ZooBank identifier urn:lsid:zoobank.org:act:86DCD029-9506-4ADA-A6BF-214A364F1EF2 = *Caenorhabditis sp. 72* (temporary number)

**Type material:** The type isolate by present designation is isofemale culture QG4708, cryopreserved as a living stock at the *Caenorhabditis* Genetics Center, Minneapolis, MN. Derived from a single female collected in Kitti, Pohnpei, Federated States of Micronesia (coordinates N 6.8652, E 158.173, elevation 167 m, 6 December 2023. The holotype is deposited at the *Caenorhabditis* Genetics Center. **Etymology:** Losolos is a Pohnpeian word meaning lush, in reference to the type locality.

**Diagnosis:** The species is diagnosed and delineated by fertile crosses with type isolate QG4708 in both directions, yielding fertile male and female offspring that are interfertile with one another and with both parental isolates in both directions. The species differs in its ITS2 DNA sequence from all other named species, including those listed in Félix *et al*. 2014; Huang *et al*. 2014; Ferrari *et al*. 2017; Slos *et al*. 2017; Kanzaki *et al*. 2018; Crombie *et al*. 2019; Stevens *et al*. 2019; Dayi *et al*. 2021; Sloat *et al*. 2022; Devi *et al*. 2025; and all other species described in this paper. The phylogeny inferred from transcriptome sequences places *C. losolos* on a long branch with no close relatives, sister to a clade that includes *C. japonica, C. imperialis*, and *C. spp. 25* and *49,* and new Pohnpeian species *C. mwetiwel, C. nansapw,* and *C ileile*. Experimental crosses with the other species described in this paper (including *C. mwetiwel, C. nansapw,* and *C ileile*) did not produce fertile progeny.

**Type locality:** The type isolate was recovered from a rotting *Citrus aurantifolia* collected from an agroforest plot at Pehleng, Kitti, on 6 December 2023. Overnight incubation in a Baermann funnel yielded approximately 20 worms, of which all but one were *Caenorhabditis* males and females. Four isofemale lines were established by picking single females on 7 December 2023 and transported to New York. These four isolates (QG4708, QG5173, QG5234, and QG5242) are all *C. losolos*. Eight additional isolates of this species were recovered from three other agroforest samples: a rotting nihn fig (*Ficus tinctoria*) in Kitti, and a rotting Hibiscus flower and a rotting cherry (*Muntingia calabura*) in U. **Morphology notes:** The male tail has the typical characteristics of Elegans supergroup species, including a heart-shaped fan with a serrated edge and terminal notch (Figure A1). Rays 1, 5, and 7 open dorsally. Ray 3 is equally distant from rays 2 and 4. Ray 4 is shorter and skinnier than ray 5. Spicules are long and slender with pointy tips, and the precloacal lip has the characteristic hook shape.

**Reproduction**: This species has separate males and females. Mating is in the parallel position on plates and females are oviparous.

***Caenorhabditis mwetiwel* Rockman, Tintori, Nguyen, & Yomai *sp. nov*.**

ZooBank identifier urn:lsid:zoobank.org:act:A4640623-3376-45F9-81BC-EE73C01135FF = *Caenorhabditis sp. 73* (temporary number)

**Type material:** The type isolate by present designation is isofemale culture QG4797, cryopreserved as a living stock at the *Caenorhabditis* Genetics Center, Minneapolis, MN. Derived from a single female collected in Nett, Pohnpei, Federated States of Micronesia (coordinates N 6.9582, E 158.2096, elevation 34 m, 13 December 2023. The holotype is deposited at the *Caenorhabditis* Genetics Center.

**Etymology:** Mwetiwel is a Pohnpeian word meaning garden, a reference to the type locality. **Diagnosis:** This species reproduces with separate males and females. The species is diagnosed and delineated by fertile crosses with type isolate QG4797 in both directions, yielding fertile male and female offspring that are interfertile with one another and with both parental isolates in both directions. The species differs in its ITS2 DNA sequence from all other named species, including those listed in Félix *et al*. 2014; Huang *et al*. 2014; Ferrari *et al*. 2017; Slos *et al*. 2017; Kanzaki *et al*. 2018; Crombie *et al*. 2019; Stevens *et al*. 2019; Dayi *et al*. 2021; Sloat *et al*. 2022; Devi *et al*. 2025; and all other species described in this paper. The phylogeny inferred from transcriptome sequences places *C. mwetiwel* as sister to a clade of *C. imperialis, C. nansapw,* and *C ileile*. Reciprocal experimental crosses with these species did not produce fertile hybrids, though all crosses but one produced viable, infertile F_1_ adults (Table S5). Crosses between *C. mwetiwel* males and *C. imperialis* females resulted in dead embryos only. The clade that includes these species and *C. mwetiwel* is in turn sister to the clade of *C. sp. 25* and *C. sp. 49*. Reciprocal experimental crosses with these species produced dead embryos, plus a small number of arrested L1 larvae in crosses of *C. mwetiwel* females and *C. sp. 25* males.

**Type locality:** The type isolate was recovered from an unidentified substrate collected from the ground in the Pohnpei Botanical Garden in Nett on 13 December 2023. Overnight incubation in a Baermann funnel yielded thousands of worms. Five isofemale lines were established by picking single females on 15 December 2023. These five isolates (QG4797, QG5291, QG5306, QG5308, and QG5338) are all *C. mwetiwel*. Six additional isolates of this species were recovered from two other samples, both from an agroforest plot in U: a rotting sawa stem (taro, *Colocasia esculenta*) and a rotting banana flower.

**Morphology notes:** The male tail has the typical characteristics of Elegans supergroup species, including a heart-shaped fan with a serrated edge and terminal notch (Figure A1). Rays 1, 5, and 7 open dorsally. Ray 3 is equally distant from rays 2 and 4. Ray 4 is shorter and skinnier than ray 5. Spicules are long and slender with pointy tips, and the precloacal lip has the characteristic hook shape.

**Reproduction**: This species has separate males and females. Mating is in the parallel position on plates and females are oviparous.

***Caenorhabditis nansapw* Rockman, Tintori, Nguyen, & Yomai *sp. nov*.**

ZooBank identifier urn:lsid:zoobank.org:act:1693D8EC-B243-4B2B-A3EA-BF128572BF90 *= Caenorhabditis sp. 74* (temporary number)

**Type material:** The type isolate by present designation is isofemale culture QG4644, cryopreserved as a living stock at the *Caenorhabditis* Genetics Center, Minneapolis, MN. Derived from a single female collected in Kitti, Pohnpei, Federated States of Micronesia (coordinates N 6.8632, E 158.1765, elevation 253 m, 6 December 2023. The holotype is deposited at the *Caenorhabditis* Genetics Center.

**Etymology:** Nansapw is a Pohnpeian word for cultivated land, a reference to the type locality. **Diagnosis:** This species reproduces with separate males and females. The species is diagnosed and delineated by fertile crosses with type isolate QG4644 in both directions, yielding fertile male and female offspring that are interfertile with one another and with both parental isolates in both directions. The species differs in its ITS2 DNA sequence from all other named species, including those listed in Félix *et al*. 2014; Huang *et al*. 2014; Ferrari *et al*. 2017; Slos *et al*. 2017; Kanzaki *et al*. 2018; Crombie *et al*. 2019; Stevens *et al*. 2019; Dayi *et al*. 2021; Sloat *et al*. 2022; Devi *et al*. 2025; and all other species described in this paper. The phylogeny inferred from transcriptome sequences places *C. nansapw* as sister to *C. ileile*. Reciprocal experimental crosses with *C. ileile* produced viable F_1_s in both directions, but these F_1_s did not produce any F_2_s. Reciprocal crosses between each class of F_1_ and each parental isolate produced zero or few embryos, which failed to hatch, in every case but one: F_1_ females from the cross of *C. nansapw* females and *C. ileile* males, when crossed to *C. nansapw* males, produce a small number of offspring, some of which developed to adulthood. The clade of *C. nansapw* and *C. ileile* is sister to *C. imperialis,* with *C. mwetiwel* as the next outgroup.

Reciprocal experimental crosses between *C. nansapw* and these other species produced infertile adults in all cases but one: crosses between *C. nansapw* females and *C. imperialis* males produced dead embryos only. Sister to this clade of four species is the clade of *C. sp. 25* and *C. sp. 49.* Reciprocal experimental crosses between *C. nansapw* and these species produced dead embryos, plus a small number of arrested L1 larvae in crosses of *C. nansapw* females and *C. sp. 25* males (Table S5).

**Type locality:** The type isolate was recovered from a rotting breadfruit collected from an agroforest plot at Pehleng, Kitti, on 6 December 2023. Overnight incubation in a Baermann funnel yielded approximately thousands of worms. Four isofemale lines were established by picking single females on 7 December 2023. These four isolates (QG4644, QG5287, QG5292, and QG5355) are all *C. nansapw*. Seventeen additional isolates of this species were recovered from five other samples: a rotting nihn fig (*Ficus tinctoria*) and two samples of rotting false durian in disturbed forest in Sokehs, an unidentified substrate at the Pohnpei Botanical Garden in Nett, and rotting breadfruit at an agroforest plot in U.

**Morphology notes:** The male tail has the typical characteristics of Elegans supergroup species, including a heart-shaped fan with a serrated edge and terminal notch (Figure A1). Rays 1, 5, and 7 open dorsally. Ray 3 is equally distant from rays 2 and 4. Ray 4 is shorter and skinnier than ray 5. Spicules are long and slender with pointy tips, and the precloacal lip has the characteristic hook shape.

**Reproduction**: This species has separate males and females. Mating is in the parallel position on plates and females are oviparous.

***Caenorhabditis ileile* Rockman, Tintori, Nguyen, & Yomai *sp. nov*.**

ZooBank identifier urn:lsid:zoobank.org:act:61155085-3F94-4703-B045-ED9378A55FDC *= Caenorhabditis sp. 75*

**Type material:** The type isolate by present designation is isofemale culture QG4848, cryopreserved as a living stock at the Caenorhabditis Genetics Center, Minneapolis, MN. Derived from a single female collected in Kitti, Pohnpei, Federated States of Micronesia (coordinates N 6.857742, E 158.2156, elevation 671 m, 8 December 2023. The holotype is deposited at the *Caenorhabditis* Genetics Center.

**Etymology:** Ileile is a Pohnpeian word meaning very high, a reference to the type locality. **Diagnosis:** This species reproduces with separate males and females. The species is diagnosed and delineated by fertile crosses with type isolate QG4848 in both directions, yielding fertile male and female offspring that are interfertile with one another and with both parental isolates in both directions. The species differs in its ITS2 DNA sequence from all other named species, including those listed in Félix *et al*. 2014; Huang *et al*. 2014; Ferrari *et al*. 2017; Slos *et al*. 2017; Kanzaki *et al*. 2018; Crombie *et al*. 2019; Stevens *et al*. 2019; Dayi *et al*. 2021; Sloat *et al*. 2022; Devi *et al*. 2025; and all other species described in this paper. The phylogeny inferred from transcriptome sequences places *C. ileile* as sister to *C. nansapw*. Reciprocal experimental crosses with *C. nansapw* produced viable F_1_s in both directions, but these F_1_s did not produce any F_2_s. Reciprocal crosses between each class of F_1_ and each parental isolate produced zero or few embryos, which failed to hatch, in every case but one: F_1_ females from the cross of *C. nansapw* females and *C. ileile* males, when crossed to *C. nansapw* males, produce a small number of offspring, some of which developed to adulthood. The clade of *C. nansapw* and *C. ileile* is sister to *C. imperialis,* with *C. mwetiwel* as the next outgroup. Reciprocal experimental crosses between *C. ileile* and *C. imperialis* produced dead embryos. Reciprocal experimental crosses between *C. ileile* and *C. mwetiwel* produced infertile F_1_ adults. Sister to this clade of four species is the clade of *C. sp. 25* and *C. sp. 49.* Reciprocal experimental crosses between *C. ileile* and these species produced dead embryos, plus a small number of arrested L1 larvae in crosses of *C. ileile* females and *C. sp. 49* males (Table S5).

**Type locality:** The type isolate was recovered from rotting kotop (*Clinostigma ponapensis*) palm fruits collected from cloudforest along the ridge above Enipein, Kitti, on 8 December 2023. Overnight incubation in a Baermann funnel yielded many male and female *Caenorhabditis* larvae. Four isofemale lines were established by picking single females on 9 December 2023. One of these is QG4848, the type isolate of *C. ileile*. The other three isolates (QG4862, QG4729, and QG4884) are *C. pwilidak*. *C. ileile* is known only from this single isolate.

**Morphology notes:** The male tail has the typical characteristics of Elegans supergroup species, including a heart-shaped fan with a serrated edge and terminal notch (Figure A1). Rays 1, 5, and 7 open dorsally. Ray 3 is equally distant from rays 2 and 4. Ray 4 is shorter and skinnier than ray 5. Spicules are long and slender with pointy tips, and the precloacal lip has the characteristic hook shape.

**Reproduction**: This species has separate males and females. Mating is in the parallel position on plates and females are oviparous.

***Caenorhabditis pwilidak* Rockman, Tintori, Nguyen, & Yomai *sp. nov*.**

ZooBank identifier urn:lsid:zoobank.org:act:D207B048-A366-4422-904B-A169AE0D5244 *= Caenorhabditis sp. 76*

**Type material:** The type isolate by present designation is isofemale culture QG4628, cryopreserved as a living stock at the *Caenorhabditis* Genetics Center, Minneapolis, MN. Derived from a single female collected in Kitti, Pohnpei, Federated States of Micronesia (coordinates N 6.906566, E 158.1817, elevation 541 m, 6 December 2023. The holotype is deposited at the *Caenorhabditis* Genetics Center.

**Etymology:** Pwilidak is the Pohnpeian word for native, as in reference to a native of Pohnpei. **Diagnosis:** This species reproduces with separate males and females. The species is diagnosed and delineated by fertile crosses with type isolate QG4628 in both directions, yielding fertile male and female offspring that are interfertile with one another and with both parental isolates in both directions. The species differs in its ITS2 DNA sequence from all other named species, including those listed in Félix *et al*. 2014; Huang *et al*. 2014; Ferrari *et al*. 2017; Slos *et al*. 2017; Kanzaki *et al*. 2018; Crombie *et al*. 2019; Stevens *et al*. 2019; Dayi *et al*. 2021; Sloat *et al*. 2022; Devi *et al*. 2025; and all other species described in this paper. The phylogeny inferred from transcriptome sequences places *C. pwilidak* as distant sister to the clade of *C. kamaaina* and *C. oiwi*. Reciprocal experimental crosses between *C. pwilidak* and each of these species resulted in dead F_1_ embryos in each case (Table S5).

**Type locality:** The type isolate was recovered from a rotting pwuhr (*Fagraea berteroana*) fruit collected from cloudforest along the ridge above Pehleng, Kitti, on 6 December 2023. Overnight incubation in a Baermann funnel yielded 20-30 *Caenorhabditis* larvae. Five isofemale lines were established by picking single females on 9 and 10 December 2023. All five isolates (QG4628, QG4643, QG5307, QG5325, and QG5332) are *C. pwilidak*. We collected an additional 801 isolates of *C. pwilidak*. These include 425 isolates from 71 substrates collected in three 1-m^2^ grids of ninh (*Ficus tinctoria*) figs at Sokehs Ridge, and 381 collected from 101 samples representing 17 diverse substrate types in Kitti, Madolenhimw, Sokehs, and U.

**Morphology notes:** The male tail has the typical characteristics of Elegans supergroup species, including a heart-shaped fan with a serrated edge and terminal notch (Figure A1). Rays 1, 5, and 7 open dorsally. Ray 3 is equally distant from rays 2 and 4. Ray 4 is the same length and thickness as ray 5, which is different from its closest relatives *C. kamaaina* and *C. oiwi* (Crombie *et al*. 2019). Spicules are long and slender with pointy tips, and the precloacal lip has the characteristic hook shape.

**Reproduction**: This species has separate males and females. Mating is in the parallel position on plates and females are oviparous.

**Figure A1.**
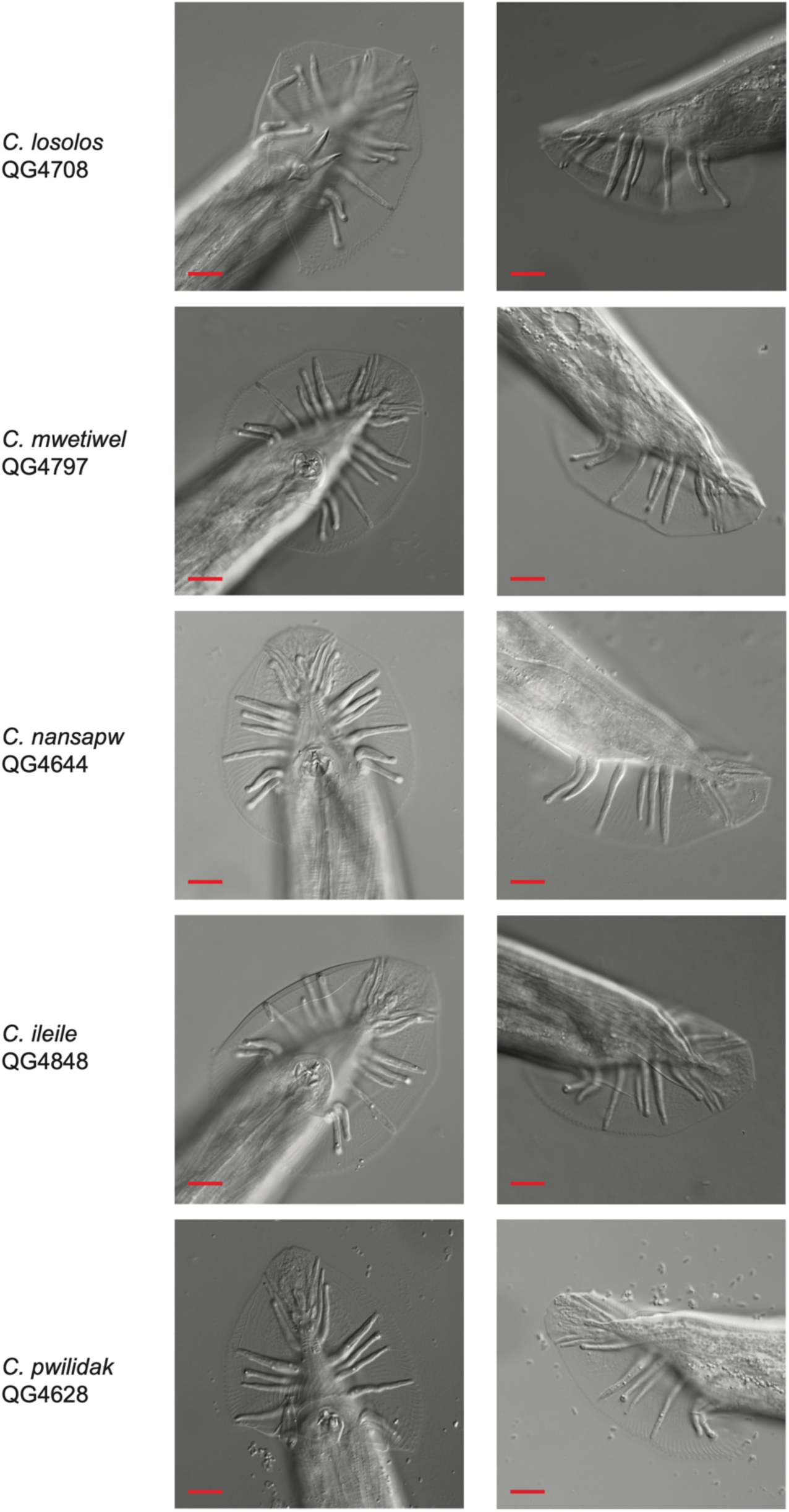
Ventral and lateral views of the tails of adult males, one day post-L4, for the type strains of each of five new species of *Caenorhabditis*. Scale bars are 10 µm.

