## Supplementary material for "*Caenorhabditis* diversity on Pohnpei, Micronesia, provides evidence that the Elegans Supergroup has its roots in the Americas and diversified in the Pacific en route to Asia": FileS1.StatisticsOfPatchColonization.R

```
################################################################
# File S1. Scripts for colonization statistics
################################################################

# Numerically find the likelihood surface for lambda_C and lambda_D, given 7 1-worm samples out of 101 ≥1-worm samples
# This result is plotted in Figure 4B.


# Data:
n <- 101
k <- 7

# Parameter values to consider:
lambda_C <- seq(0.01, 3, 0.01) # 0.01 to 3 colonizations per patch 
lambda_D <- seq(0.1, 10, 0.01) # 0.1 to 10 dauers per colonization

# conversion of lambda_D to estimated founder dauer number (i.e., from Poisson mean to zero-truncated Poisson mean)
nd_hat <- lambda_D/(1-exp(-lambda_D))

# Set up resuts matrix
likelihood <- matrix(nrow = length(lambda_D), ncol = length(lambda_C))

# Calculate likelihoods
for(j in 1:length(lambda_C)){
	ncol <- lambda_C[j]
		for(q in 1:length(lambda_D)){
			nworms <- lambda_D[q]
			p <- ncol*nworms/((exp(ncol)-1)*(exp(nworms)-1))
			likelihood[q,j] <- log(dbinom(k,n,p)	)		
		}	
}

max(likelihood)
# [1] -1.867939


# Plot Likelihood surface
# Colored region corresponds to 2-parameter 95% CI (a log likelihood drop of qchisq(0.95,2)/2))
image(x = nd_hat, y = lambda_C, z = likelihood, zlim = c(max(likelihood) - qchisq(0.95,2)/2, max(likelihood)), col = hcl.colors(12, "YlOrRd", rev = T), xlim = c(1,10))
contour(x = nd_hat, y = lambda_C, z =likelihood, add =T, nlevels = 40)

# Find the maximum likelihood estimate of lambda_D at each value of lambda_C
estimate <- NULL
ml_D <- NULL
for(j in 1:length(lambda_C)){
	ncol <- lambda_C[j]
		maxD <- optimize(function(x){log(dbinom(k,n,(dpois(1, ncol)/(1-dpois(0, ncol))) * dpois(1,x)/(1-dpois(0,x))))}, interval = c(0,50), maximum = T)			
	estimate[j] <- maxD $maximum
	ml_D[j] <- maxD $objective
}	
# This shows that there is a ridge, with the maximum likelihood = -1.867939. 
# Convert the lambda_D estimates to the zero-truncated nd (number of dauers)
nd_est_max <- estimate/(1-exp(-estimate))

# Add to the plot
points(lambda_C ~ nd_est_max, ty = "l", lwd = 5, col = "red")

# Describe the ridge with a quadratic
summary(lm(nd_est_max ~ lambda_C + I(lambda_C^2)))$coef
#                 Estimate   Std. Error   t value     Pr(>|t|)
# (Intercept)    4.17106064 0.0004661486 8947.9198  0.00000e+00
# lambda_C      -0.63204357 0.0007151505 -883.7910  0.00000e+00
# I(lambda_C^2) -0.04218941 0.0002300950 -183.3565 1.27412e-307

################################################################

# Estimate the number of colonizations per patch, using 0-worm patches
# From the 400-sample survey
# This result contributes to Figure 4C

n_patches <- 400
n_Cpwilidak <- 101

# Estimate if we assume all 400 patches are available for colonization
lambda_C_total <- optimize(function(y){log(dbinom(n_patches - n_Cpwilidak, n_patches, dpois(0,y)))}, interval = c(0,2), maximum = T)
lambda_C_total
# $maximum
# [1] 0.2910194
# $objective
# [1] -3.081884

# Plot the likelihood surface
lseq <- seq(0.1, 0.5, 0.001)
lnL <- rep(0,length(lseq))
for(i in 1:length(lseq)){
	lnL[i] <- log(dbinom(n_patches - n_Cpwilidak, n_patches, dpois(0,lseq[i])))
}
plot(lnL ~ lseq)
abline(v = lambda_C_total$maximum, col = "red")
# Find the 95% CI via 1.92-log likelihood drop (using 1df Chisq)
lambda_C_total$objective - qchisq(0.95,1)/2
# [1] -5.001884
abline(h = lambda_C_total$objective - qchisq(0.95,1)/2, lty = 2)
# CI: 0.237 - 0.352

# One these assumptions, how many C. pwilidak-colonized patches received exactly one colonization event?
dpois(1, lambda_C_total$maximum)/(1-dpois(0, lambda_C_total$maximum))
# [1] 0.8615381

# And therefore 1-0.8615381 = 0.1384619 received more than one. 

################################################################

# Estimate dauer number nd, conditional on lambda_C = 0.2910194
# This result contributes to Figure 4C

fixed_lambda_C <- lambda_C_total$maximum

# Find the maximum likelihood lambda_D
MLfixedC <- optimize(function(x){log(dbinom(k,n,(dpois(1, fixed_lambda_C)/(1-dpois(0, fixed_lambda_C))) * dpois(1,x)/(1-dpois(0,x))))}, interval = c(0,50), maximum = T)
lambda_D_hat_fixedC <- MLfixedC$maximum
# Convert that value to an estimate of the number of founders, F_hat			
founders <- lambda_D_hat_fixedC/(1-exp(-lambda_D_hat_fixedC))
founders
# [1] 3.982549

# Plot the likelihood surface
lseq <- seq(1.5,7.5,.01)
lnL <- rep(0,length(lseq))
for(i in 1:length(lseq)){
	lnL[i] <- log(dbinom(k,n,(dpois(1, fixed_lambda_C)/(1-dpois(0, fixed_lambda_C))) * dpois(1,lseq[i])/(1-dpois(0,lseq[i]))))
}
founderseq <- lseq/(1-exp(-lseq))
plot(lnL ~ founderseq)

abline(v = founders, col = 2)
# Find the 95% CI via 1.92-log likelihood drop (using 1df Chisq)
MLfixedC$objective - qchisq(0.95,1)/2
# [1] -3.788668
abline(h = MLfixedC$objective - qchisq(0.95,1)/2, lty = 2)
# CI = 3.20 - 5.00 in founder count
# CI = 3.05 - 4.96 in lambda_D 


################################################################

# Estimate founder number for successful patches 
# Patches with one worm or worms of only one sex don't proliferate.
# And proliferative patches may have been founded my multiple colonization events
# Here are simulations that use lambda_C and lambda_D 


# Case with 50:50 sex ratio
# This result is plotted in Figure 4C. 

# Simulate colonizations for many patches
patches <- rpois(1e7, fixed_lambda_C)
# Consider only those patches that received worms 
patches <- patches[patches >0]
# For each colonization, add a poisson-distributed number of worms
for(i in 1:length(patches)){
	dauercount <- rpois(50, lambda_D_hat_fixedC)
	dauercount <- dauercount[dauercount > 0] # Zero-trucate the distribution
dauers[i] <- sum(sample(dauercount, patches[i])) 
}
# Now we have a vector of the number of founders for each colonized patch, including the possibility of founders from multiple colonizations of a patch.
# Now test whether each patch has both males and females
# sex ratio specified by male frequency m
m <- 0.5
males <- NULL
for(j in 1:length(dauers)){
	males[j] <- rbinom(1, dauers[j], m)
}
# Find the patches with only males or only females
allmales <- dauers -males == 0
allfemales <- dauers-males == dauers
# The successes are whatever is left:
successes <- dauers[allmales ==F & allfemales ==F]
successes <- successes[1:1e6]
summary(successes)
hist(successes, br = seq(1.5, max(successes)+0.5, 1))

################################################################

# Now consider a biased sex ratio

m <- 0.2
males <- NULL
for(j in 1:length(dauers)){
	males[j] <- rbinom(1, dauers[j], m)
}
# Find the patches with only males or only females
allmales <- dauers -males == 0
allfemales <- dauers-males == dauers
# The successes are whatever is left:
successes <- dauers[allmales ==F & allfemales ==F]
successes <- successes[1:1e6]
summary(successes)
hist(successes, br = seq(1.5, max(successes)+0.5, 1))

################################################################

# Now find the CIs for these estimates:
# Use 3.05 and 4.96 as the lower and upper bounds of the CI estimated for lambda_D when lambda_C = 0.2910194. That is, we're only considering error in lambda_D here. 

# Low end of the CI
patches <- rpois(1e7, fixed_lambda_C)
patches <- patches[patches >0]
for(i in 1:length(patches)){
	dauercount <- rpois(50, 3.05) ##### <-------- This is where we change the dauers per patch number to estimate the low end of the CI
	dauercount <- dauercount[dauercount > 0]
dauers[i] <- sum(sample(dauercount, patches[i])) 
}
m <- 0.5
males <- NULL
for(j in 1:length(dauers)){
	males[j] <- rbinom(1, dauers[j], m)
}
allmales <- dauers -males == 0
allfemales <- dauers-males == dauers
successes <- dauers[allmales ==F & allfemales ==F]
successes <- successes[1:1e6]
summary(successes)
#   Min. 1st Qu.  Median    Mean 3rd Qu.    Max. 
#  2.000   3.000   4.000   4.473   5.000  27.000 
hist(successes, br = seq(1.5, max(successes)+0.5, 1))

# And now the high end of the CI
patches <- rpois(1e7, fixed_lambda_C)
patches <- patches[patches >0]
for(i in 1:length(patches)){
	dauercount <- rpois(50, 4.96) ##### <-------- This is where we change the dauers per patch number to estimate the high end of the CI
	dauercount <- dauercount[dauercount > 0]
dauers[i] <- sum(sample(dauercount, patches[i])) 
}
m <- 0.5
males <- NULL
for(j in 1:length(dauers)){
	males[j] <- rbinom(1, dauers[j], m)
}
allmales <- dauers -males == 0
allfemales <- dauers-males == dauers
successes <- dauers[allmales ==F & allfemales ==F]
successes <- successes[1:1e6]
summary(successes)
#   Min. 1st Qu.  Median    Mean 3rd Qu.    Max. 
#  2.000   4.000   6.000   6.227   8.000  37.000 
hist(successes, br = seq(1.5, max(successes)+0.5, 1))

# And low and high CI boundaries when m = 0.2:
patches <- rpois(1e7, fixed_lambda_C)
patches <- patches[patches >0]
for(i in 1:length(patches)){
	dauercount <- rpois(50, 3.05) ##### <-------- This is where we change the dauers per patch number to estimate the low end of the CI
	dauercount <- dauercount[dauercount > 0]
dauers[i] <- sum(sample(dauercount, patches[i])) 
}
m <- 0.2. ##### <-------- This is where we change the sex ratio
males <- NULL
for(j in 1:length(dauers)){
	males[j] <- rbinom(1, dauers[j], m)
}
allmales <- dauers -males == 0
allfemales <- dauers-males == dauers
successes <- dauers[allmales ==F & allfemales ==F]
successes <- successes[1:1e6]
summary(successes)
#   Min. 1st Qu.  Median    Mean 3rd Qu.    Max. 
#  2.000   3.000   4.000   4.687   6.000  27.000 
hist(successes, br = seq(1.5, max(successes)+0.5, 1))

patches <- rpois(1e7, fixed_lambda_C)
patches <- patches[patches >0]
for(i in 1:length(patches)){
	dauercount <- rpois(50, 4.96) ##### <-------- This is where we change the dauers per patch number to estimate the high end of the CI
	dauercount <- dauercount[dauercount > 0]
dauers[i] <- sum(sample(dauercount, patches[i])) 
}
m <- 0.2.  ####<-------- This is where we change the sex ratio
males <- NULL
for(j in 1:length(dauers)){
	males[j] <- rbinom(1, dauers[j], m)
}
allmales <- dauers -males == 0
allfemales <- dauers-males == dauers
successes <- dauers[allmales ==F & allfemales ==F]
successes <- successes[1:1e6]
summary(successes)
#   Min. 1st Qu.  Median    Mean 3rd Qu.    Max. 
#  2.000   4.000   6.000   6.581   8.000  37.000 
hist(successes, br = seq(1.5, max(successes)+0.5, 1))


################################################################

# Statistics for the cooccurrence of C. pwilidak and C. briggsae at Sokehs Ridge
# These analyses contribute to the Discussion section headed "Factors maintaining species diversity."

# Estimate the number of colonizations per patch, using 0-worm patches from Sokehs Ridge
# Number of figs with data from the spatial array experiment:
n_figs <- 74

# Of the n=74 patches, how many contained zero of each species?
k_Cpwilidak <- 22
k_Cbriggsae <- 60

#####

# Estimate lambda_C for C. pwilidak:
Cpw_lambdaC <- optimize(function(y){log(dbinom(k_Cpwilidak, n_figs, dpois(0,y)))}, interval = c(0,10), maximum = T)
# lambda_C estimate:
Cpw_lambdaC
# $maximum
# [1] 1.213022

# log likelihood: 
# $objective
# [1] -2.292313

# Plot the likelihood surface:
lseq <- seq(0.5, 2.4, 0.001)
lnL <- rep(0,length(lseq))
for(i in 1:length(lseq)){
	lnL[i] <- log(dbinom(k_Cpwilidak, n_figs, dpois(0,lseq[i])))
}
plot(lnL ~ lseq)

# Log likelihood at 1.92-log drop:
optimize(function(y){log(dbinom(k_Cpwilidak, n_figs, dpois(0,y)))}, interval = c(0,10), maximum = T)$obj-1.92

abline(v = 1.213022, col =2)
abline(h = -4.212313, lty = 2)

# How many C. pwilidak-colonized figs received more than one colonization by C. pwilidak?
1-dpois(1, 1.213022)/(1-dpois(0,1.213022))
# [1] 0.4867979

#####

# Estimate lambda_C for C. briggsae:
Cbr_lambdaC <- optimize(function(y){log(dbinom(k_Cbriggsae, n_figs, dpois(0,y)))}, interval = c(0,10), maximum = T)
# lambda_C estimate:
Cbr_lambdaC
# $maximum
# [1] 0.2097119

# log likelihood: 
# $objective
# [1] -2.139821

# Plot the likelihood surface:
lseq <- seq(0.05, 0.5, 0.001)
lnL <- rep(0,length(lseq))
for(i in 1:length(lseq)){
	lnL[i] <- log(dbinom(k_Cbriggsae, n_figs, dpois(0,lseq[i])))
}
plot(lnL ~ lseq)

optimize(function(y){log(dbinom(k_Cbriggsae, n_figs, dpois(0,y)))}, interval = c(0,10), maximum = T)$obj - 1.92
# [1] -4.059821

abline(v = 0.2097119, col =2)
abline(h = -4.059821, lty = 2)

# How many C. briggsae-colonized figs received more than one colonization by C. briggsae?
1-dpois(c(1), 0.2097119)/(1-dpois(0,0.2097119))
# [1] 0.1011937

#######

# Modeling Ives' 1991 J statistic
# As treated in the Discussion Section

Jp <- NULL	# J for C. pwilidak; Ives Eq 1
Jb <- NULL	# J for C. briggsae; Ives Eq 1
Cpb <- NULL	# C between pwilidak and briggsae; Ives Eq 2
nsims <- 1e6 # simulate the Sokehs Ridge experiment q times

for(q in 1:nsims){
	L <- 74 # number of patches observed at Sokehs Ridge
	
	# C. pwilidak patch colonization
	Lcolp <- rpois(74, Cpw_lambdaC$maximum) # colonizations per patch
	nisCp <- rep(0,length(Lcolp))
	for(i in 1:length(Lcolp)){ # This look adds colonists for each colonization, drawing from a zero-truncated Poisson distribution
		dauercount <- rpois(50, lambda_D_hat_fixedC)
		dauercount <- dauercount[dauercount > 0]
	nisCp[i] <- sum(sample(dauercount, Lcolp[i])) 
	} # nisC is a vector of observations of founder number for all 74 patches	
	Np <- sum(nisCp)/L # mean founders per patch
	Jp[q] <- (sum(nisCp*(nisCp-1)/(Np*L))-Np)/Np # equation 1 from Ives 1999
	
	# C. briggsae patch colonization, exactly as for C. pwilidak but with different colonization rate
	Lcolb <- rpois(74, Cbr_lambdaC$maximum) 
	nisCb <- rep(0,length(Lcolb))
	for(i in 1:length(Lcolb)){
		dauercount <- rpois(50, lambda_D_hat_fixedC)
		dauercount <- dauercount[dauercount > 0]
	nisCb[i] <- sum(sample(dauercount, Lcolb[i])) 
	}
	Nb <- sum(nisCb)/L
	Jb[q] <- (sum(nisCb*(nisCb-1)/(Nb*L))-Nb)/Nb # equation 1 from Ives 1999

	# Interspecific aggregation
	Cpb[q] <- (sum(nisCp*nisCb/(Np*L))-Nb)/Nb
}

summary(Jp)
#   Min. 1st Qu.  Median    Mean 3rd Qu.    Max. 
# 0.2075  0.6668  0.7809  0.8007  0.9126  2.6075 

summary(Jb)
#   Min. 1st Qu.  Median    Mean 3rd Qu.    Max.    NA's 
#  1.143   3.638   4.476   4.836   5.605  58.200       1 
# One of the 1-million simulations had zero colonizations by C. briggsae, yielding NA

summary(Cpb)
#     Min.    1st Qu.     Median       Mean    3rd Qu.       Max.       NA's 
#-1.0000000 -0.2001055 -0.0167177  0.0000551  0.1804298  2.3426191          1 

# Apply Ives' 1991 equation 11 to estimate the reduction in interspecific competition
summary(1 - ((Cpb+1)^2)/((Jp+1)*(Jb+1)))
#   Min. 1st Qu.  Median    Mean 3rd Qu.    Max.    NA's 
# 0.3543  0.8513  0.9003  0.8884  0.9382  1.0000       1 
```
